# Decadal tracking reveals species-specific limits to coral thermal acclimatization

**DOI:** 10.64898/2026.07.15.738835

**Authors:** Jill Ashey, Kristen T. Brown, Marcelina P. Martynek, Benjamin H. Glass, Conall McNicholl, Crawford Drury, Katie L. Barott

**Affiliations:** Department of Biology, University of Pennsylvania, Philadelphia, PA USA; Global Science and Technology, Inc., Greenbelt, MD USA; U.S. National Oceanic and Atmospheric Administration, National Environmental Satellite Data and Information Service, Center for Satellite Applications and Research, Coral Reef Watch, MD USA; Department of Biology, Boston University, Boston, MA USA; Hawaiʻi Institute of Marine Biology, University of Hawaiʻi, Kāneʻohe, HI USA

## Abstract

Mass coral bleaching events driven by marine heatwaves are increasing in frequency and severity, yet the long-term recovery trajectories of surviving corals remain poorly understood. Here, leveraging a cohort of individual coral colonies with a decade of tracked environmental and bleaching history, we captured structural shifts in the coral thermal performance landscape. Specifically, we quantified thermal performance curves (TPCs) for photosynthesis and calcification in bleaching-resistant and bleaching-susceptible colonies of two ecologically dominant reef-building corals (*Montipora capitata* and *Porites compressa*) at four and six years following the 2019 marine heatwave in Kāneʻohe Bay, Hawaiʻi (2023 and 2025, respectively). Coral thermal performance shifted substantially between 2023 and 2025, and these shifts differed between species, bleaching phenotypes, and traits. In *P. compressa*, photosynthetic thermal optimum (T_opt_) shifted downward over time by 1.6°C across both phenotypes, suggesting recalibration toward prevailing conditions at the potential cost of future heat tolerance. In *M. capitata*, photosynthetic performance was lower in bleaching-susceptible corals in 2023 but converged by 2025, suggesting susceptible colonies recovered. In contrast, T_opt_ of photosynthesis remained persistently higher in bleaching-resistant colonies, likely reflecting established symbiont communities. Critically, photosynthesis and calcification did not recover in parallel. Calcification TPCs for both phenotypes of *M. capitata* were stable across both timepoints, whereas calcification TPCs in *P. compressa* continued to change through 2025. These findings demonstrate that coral thermal performance is not static following heatwaves but continues to be reshaped over multi-year recovery periods, and that species-specific traits and strategies can fundamentally constrain the pace and coupling of physiological recovery. Furthermore, elevated thermal tolerance acquired through a heatwave can erode during prolonged periods of ambient temperatures. As recurrent bleaching events shorten recovery windows, understanding these dynamic physiological trajectories are essential for understanding and forecasting reef futures.

## 1 | Introduction

Coral reefs are among the most biodiverse and economically important ecosystems on Earth and have dramatically declined due to increasing sea surface temperatures (Heron et al. 2016; Spady et al. 2026). Mass coral bleaching events are driven by unprecedentedly warm sea surface temperatures that disrupt the obligate symbiosis between tropical corals and their photosynthetic algal endosymbionts (Glynn 1993), and have increased in frequency and severity over recent decades (Oliver et al. 2019; Spady et al. 2026). Under current fossil fuel emissions trajectories, stressful temperature conditions are projected to occur annually for most tropical reefs by the end of this century (Van Hooidonk et al. 2016; Spady et al. 2026), leaving coral reefs with short recovery windows between events. While corals possess the capacity to recover physiological functions following a bleaching event (Baker et al. 2008; Brown, Lenz, et al. 2023), understanding the dynamics and limits of recovery between recurrent bleaching events is critical for predicting reef futures.

Post-bleaching recovery is a multi-dimensional process involving symbiont repopulation, energy reserve replenishment, and cellular repair, all of which underpin the return to homeostasis (Baker et al. 2008). While the recovery of physiological indicators like symbiont density and chlorophyll content can occur relatively rapidly, often within a single seasonal cycle (Rodrigues and Grottoli 2007; Schoepf et al. 2015; Wall et al. 2021; Strand et al. 2024), these visual indicators of recovery do not always reflect a full return to physiological robustness. More energetically demanding processes, such as calcification, can remain impaired for extended periods after stress. For example, skeletal growth can remain suppressed for several years following heat stress (Cantin and Lough 2014; Gold and Palumbi 2018; Levas et al. 2018; Baumann et al. 2019). Because calcification underpins the structural integrity of the reef, its long-term suppression compromises the framework that supports reef biodiversity and ecosystem services (Wild et al. 2011). While these long-term physiological legacies highlight specific functional constraints during recovery, we lack a comprehensive framework to quantify how the relationship between heat stress and physiology shifts as corals undergo multi-year recovery from marine heatwaves.

Thermal performance curves (TPCs) offer a robust quantitative framework for describing temperature-dependence of physiological rates (Sinclair et al. 2016). By measuring physiological rates across a range of temperatures and fitting these data to mechanistic models, TPCs simultaneously capture the temperature at which performance is maximized (thermal optimum, T_opt_), the rate of performance increase below the optimum (activation energy, E_a_), the rate of thermal denaturation above the optimum (deactivation energy, E_h_), and the magnitude of performance at reference and maximum temperatures (b(T_c_) and R_max_, respectively; Figure S1) (Daniel Padfield et al. 2021). This multi-parameter characterization reveals the thermal performance landscape of an organism, and has been increasingly used to assess acclimatory capacity and responses to future stress events in marine ectotherms, including corals (Daniel Padfield et al. 2015; Sinclair et al. 2016). Recent studies have applied TPC approaches to coral photosynthesis, respiration, and calcification to infer thermal limits and vulnerability to projected warming and other environmental stressors (Aichelman et al. 2019; Jurriaans and Hoogenboom 2019; Silbiger et al. 2019; Anton et al. 2020; Becker and Silbiger 2020; Jurriaans and Hoogenboom 2020; Becker et al. 2021; Jurriaans et al. 2021). However, we lack empirical data on how these acute performance curves change over multi-year post-bleaching recovery timelines. Following a severe marine heatwave, surviving colonies may acclimatize to higher temperatures and elevate their thermal thresholds, effectively shifting their T_opt_ to track increasing baseline temperatures (Banc-Prandi et al. 2022). Alternatively, corals may fail to adjust after severe thermal stress, resulting instead in a depression of maximum performance capacity (R_max_), a flattening of metabolic acceleration (E_a_), and/or an accelerated onset of functional denaturation (E_h_). Evaluating TPCs at multiple time points following marine heatwaves is therefore essential to understand the capacity for recovery.

Understanding bleaching recovery mechanisms is further complicated by the observation that corals display intra- and inter-specific variability in bleaching susceptibility. Within coral populations, individual colonies differ in their bleaching susceptibility, with some colonies bleaching severely under thermal stress while neighboring conspecifics remain visually unaffected (Matsuda et al. 2020; Innis et al. 2021; Brown, Lenz, et al. 2023). These bleaching-resistant and bleaching-susceptible phenotypes have been well-documented in Kāneʻohe Bay, Hawaiʻi, with colonies of *Montipora capitata* and *Porites compressa* having consistently different bleaching susceptibilities across multiple heatwaves (Matsuda et al. 2020; Innis et al. 2021; Brown, Lenz, et al. 2023). Importantly, surviving bleaching is not equivalent to full physiological recovery, and the absence of bleaching is not equivalent to an absence of heat stress (Innis et al. 2021; Kruse et al. 2025). In *M. capitata* populations in Kāneʻohe Bay, susceptible colonies showed only partial recovery of symbiont density and tissue biomass three years after the 2019 heatwave, whereas *P. compressa* appeared to recover most physiological metrics within two years (Brown, Lenz, et al. 2023). Whether historical bleaching phenotype predicts recovery trajectory and whether phenotypic differences in physiological thermal performance persist over a recovery period remains an open question.

Here, we address these gaps by characterizing coral photosynthetic and calcification TPCs, along with a suite of physiological traits, in bleaching-resistant and bleaching-susceptible colonies of two ecologically dominant reef-building corals (*M. capitata* and *P. compressa*). TPC experiments were conducted in 2023 and 2025, representing four and six years of recovery following the most recent marine heatwave in Kāneʻohe Bay, Hawaiʻi (2019). This heatwave was the third of a series of heatwaves that triggered mass coral bleaching in Kāneʻohe Bay, with the two prior events occurring in 2014 and 2015 (Bahr et al. 2015, 2017). Critically, this study capitalizes on a decade of continuous monitoring of these specific individuals, starting with the 2015 heatwave. By assessing the same individuals over multiple post-heatwave timepoints, we can directly attribute changes in thermal performance to colony acclimatization rather than to selective mortality (i.e., loss of weaker individuals from the population). This study included three primary objectives: (1) to evaluate the extent to which historical bleaching phenotypes exert a physiological legacy on multi-year recovery trajectories; (2) to determine how photosynthetic and calcification thermal performance change across multiple years of recovery following a marine heatwave; and (3) to evaluate whether symbiont photosynthesis and coral calcification recover in a coupled manner. This framework links coral bleaching history with temporal changes in thermal performance in the years following a marine heatwave, providing a powerful mechanistic assessment of coral performance during multi-year post-heatwave recovery.

## 2 | Methods

### 2.1 | Site description and species selection

This study was conducted at patch reef 13 (PR13; 21.4515°N, 157.7966°W), located in the southern region of Kāneʻohe Bay, Oʻahu, Hawaiʻi. The two most abundant reef-building coral species (*Montipora capitata* and *Porites compressa*) in Kāneʻohe Bay were examined. Experimental colonies were selected based on historical bleaching phenotypes (bleaching-resistant vs. bleaching-susceptible), first identified during the 2015 marine heatwave (Matsuda et al. 2020). For each species, adjacent colony pairs with consistently contrasting bleaching phenotypes (i.e., bleaching-resistant vs. bleaching-susceptible) (Innis et al. 2021; Brown, Lenz, et al. 2023) were used for this study (n=9–12 per species per phenotype per timepoint; Tables S1, S2).

### 2.2 | In situ temperature monitoring

To characterize the thermal regimes of PR13, *in situ* temperatures were recorded using high-resolution data loggers (U22-001, accuracy ±0.2°C, resolution 0.02°C, Onset Computer Corporation; n=3) deployed at 1–2.7 m depth. Loggers were deployed in April 2022, and recorded temperature at 1-hr intervals. To provide historical (i.e., 2015–2022) temperature data, temperature data from the same site (PR13) was obtained from Brown et al. 2023 (Figure S2). Missing data was supplemented with data from *in situ* water temperature sensors (YSI 44032, accuracy ±0.2°C, resolution 0.1°C, Yellow Springs Instruments) at Moku o Loʻe (Station MOKH1 1612480, NOAA), which is located ∼2 km from PR13 within the southern region of Kāneʻohe Bay. Analysis of thermal regimes was confined to March–October in order to capture the onset of seasonal warming and peak summer temperatures. Thermal stress was assessed using two localized thresholds for Kāneʻohe Bay: the maximum monthly mean (MMM; 27.3°C) and the local coral bleaching threshold (MMM + 1°C; 28.3°C) (Brown, Lenz, et al. 2023). Accumulated heat stress was calculated as Degree Heating Weeks (DHWs) following the equations in (Brown, Eyal, et al. 2023).

### 2.3 | Coral collection

Fragments (∼3–4 cm) from each colony were collected on September 21, 2023 and October 1, 2025. Prior to collections, photos of each colony were taken with a digital camera (Olympus Tough TG-4 or Canon G7x Mark II), and coral color was quantified in ImageJ as the measured red channel intensity relative to a color standard (DKC-Pro White Balance & Color Calibration Chart, DGK Color Tools) (Schindelin et al. 2012). Fragments were removed from colonies using stainless steel bone cutters and transferred to coolers (120 L) filled with ambient seawater for transit to the Hawaiʻi Institute of Marine Biology. Upon arrival, fragments were suspended from nylon fishing line in flow-through outdoor tanks supplied with ambient, sand-filtered seawater (∼50 µm) from Kāneʻohe Bay. Tanks were outfitted with 70% shade cloth to match the light conditions at the collection site (∼500 µmol m^-2^ s^-1^).

### 2.4 | Photosynthesis-irradiance curves

To determine the optimal saturating light intensity for subsequent thermal performance curve (TPC) assays, photosynthesis-irradiance (PI) curves were generated for each species, as previously described (Brown, Lenz, et al. 2023). Briefly, fragments were incubated in individual chambers of seawater, each of which was fitted with an optical oxygen (PSt7, Presens) and temperature sensors (pt1000, Presens) connected to an OXY-10 (Presens). To account for possible microbial background respiration, blank chambers containing only seawater were run in parallel for each incubation. Mixing was maintained via magnetic stir bars to prevent the formation of diffusive boundary layers. Temperature and dissolved oxygen concentrations were measured every three seconds over increasing increments of light over ∼80 mins. Light levels ranged from 150–1190 µmol m^-2^ s^-1^ in September 2023 (step size ∼175 µmol m^-2^ s^-1^; ∼10 mins step^-1^), and from 220–1570 µmol m^-2^ s^-1^ in October 2025 (step size ∼225 µmol m^-2^ s^-1^; ∼10 mins step^-1^). Light was generated by LED lights (Radion XR30 G6 Pro LED Light, EcoTech Marine), and light intensities were verified using a PAR meter at each step (MQ-150 Full Spectrum Underwater LED PAR Meter, Apogee). After measurements were completed at the maximum light levels, lights were turned off (0 µmol m^-2^ s^-1^) to measure light-enhanced dark respiration for 15 mins. Fragments were then snap-frozen in liquid nitrogen and stored at -80°C until physiological processing.

Rates of oxygen flux for each light level were determined using localized linear regressions with the LoLinR package (v0.0.9; (Olito et al. 2017)) in R (v4.5.0; (R Core Team 2024)), with rates corrected for chamber volume displaced by the fragments, blank seawater control rates, and normalized to fragment surface area using the wax-dipping method (Veal et al. 2010). To estimate photosynthetic parameters for each fragment, P-I curves were generated by fitting to the Platt model (Platt et al. 1980) using nonlinear least squares regression with the nlsLM function from the minpack.lm R package (v1.2-4; (Elzhov et al. 2023)). The model is as follows:

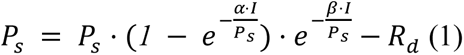

Where *P_net_* is the net photosynthetic rate (µmol O_2_ cm^-2^ h^-1^), *I* is the photosynthetic photon flux density (PPFD; µmol m^-2^ s^-1^), *P_s_* is a scaling parameter representing the theoretical maximum potential photosynthetic rate if *β* = 0, *α* is the initial slope of the curve representing photosynthetic efficiency at sub-saturating light levels, *β* is the photoinhibition parameter representing the decline in photosynthesis at high light intensities, and *R_d_* is the dark respiration rate. The saturating irradiance was derived from the model as:

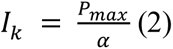

Where *I_k_* is the saturating irradiance (µmol m^-2^ s^-1^), *P_max_* is the maximum photosynthetic rate (µmol O_2_ cm^-2^ h^-1^) as derived from the Platt model, and *α* is the initial slope of the curve as derived from the Platt model.

### 2.5 | Physiological analyses

Coral tissue was removed from the skeleton by airbrushing with 50 mL of ice-cold 1x phosphate buffered saline (PBS). The resulting tissue slurry was homogenized at 20,000 rpm for 10 s using a homogenizer (Fisherbrand 850 Homogenizer, Fisher Scientific). To separate host and symbiont fractions, the homogenate was centrifuged at 4°C for 5 mins at 2,500 x *g*. The host-containing supernatant was decanted, and 1 mL aliquots were reserved for biomass quantification. The symbiont pellet was resuspended in 10 mL of 1x PBS and centrifuged again (2,500 x *g* for 5 mins at 4°C), after which the final pellet was resuspended in 900 µL of 0.1% sodium dodecyl sulfate (SDS) in 1x PBS. Symbiont concentrations were measured in triplicate on a Millipore Guava flow-cytometer (Guava easyCyte 5HT) (Krediet et al. 2015). Cells were excited with a blue laser (488 nm), and symbionts were identified by analyzing forward scatter and red fluorescence emission in GuavaSoft v3.4 with the same gating for all samples. When flow cytometry was unavailable, symbiont cells were quantified using a hemocytometer (n=6 replicate counts per sample). For biomass, 1 mL of host homogenate was dried in pre-combusted aluminum pans in an oven at 60°C for 24 hrs until a constant dry weight was achieved. Samples were then combusted in a muffle furnace at 450°C for 4–6 hrs to determine ash weight. Ash-free dry weight (AFDW) was calculated as the difference between the dry weight and ash weight. Physiological parameters were standardized to total homogenate volume and surface area, the latter of which was determined by the wax-dipping method (Veal et al. 2010)

### 2.6 | Thermal performance curve experiments

Acute thermal performance curves (TPC) were generated for net photosynthesis and light calcification across seven temperatures (18, 22, 25, 28, 31, 34, and 38°C) on September 22–26, 2023 and October 7–11, 2025. Incubations were conducted in sequence from low to high temperatures in sealed chambers filled with 50 µm filtered seawater and fitted with oxygen and temperature sensors as described above. Incubations were performed under saturating irradiance (∼730 µmol m^-2^ s^-1^ in September 2023; ∼790 µmol m^-2^ s^-1^ in October 2025), as determined by P-I curves. Corals were held at each temperature for one hour, and oxygen evolution was quantified as described above. To maintain assay temperature, seawater temperature was controlled using temperature probes and 832 Energy Bar (Apex, Neptune Systems) paired with aquarium heaters (50W: Precision Submersible Heaters, Marineland; 300W: Titanium Heater Element, Bulk Reef Supply; 500W: Orlushy Submersible Aquarium Heater) and chillers (Poafamx Aquarium Chiller).

Light calcification was measured at each temperature step using the alkalinity anomaly technique (Chisholm and Gattuso 1991). At the start of each temperature assay, baseline seawater samples were collected from the holding tank. Following the incubation period, seawater was collected from each individual chamber, including blanks. Total alkalinity was determined using an open cell titration method (SOP3b; (Dickson et al. 2007)) with a certified HCl titrant (Metrohm USA hydrochloric acid, 0.1 N). Measurements exhibited <1% error when compared against certified reference materials (Dickson Lab CRM Batches 206 and 224). Seawater changes were conducted between each temperature incubation. As described above, photosynthetic and calcification rates were adjusted for the seawater volume displaced by each coral fragment, corrected for background rates in blank chambers, and normalized to fragment surface area.

### 2.7 | TPC modeling and parameter extraction

Individual TPCs for each performance rate (net photosynthesis and calcification) were fitted to the Sharpe-Schoolfield model (Sharpe and DeMichele 1977; Schoolfield et al. 1981):

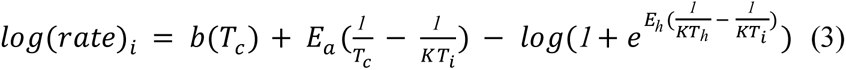

Where *i* is the individual coral fragment for each rate, *b(T_c_)* is the log rate of performance at a reference temperature (µmol O_2_ cm^-2^ h^-1^), *E_a_* is the activation energy (electron volts, eV), *T_c_* is the reference temperature at which no low or high-temperature inactivation is experienced (defined here as 300.15 K or 27°C), *K* is the Boltzmann constant (8.62 x 10^-5^ eV K^-1^), *T_i_* is the temperature in Kelvin (K), *E_h_* is the temperature-induced inactivation of enzymes past *T_h_* for each population (eV), and *T_h_* is the temperature (K) at which half the enzymes are inactivated. Models were fitted using the nls_multstart function from the nls.multstart package (v2.0.0; (Padfield et al. 2025)). From the fitted TPCs for each fragment, the following TPC metrics were extracted (Figure S1):

- T_opt_ (Acute thermal optimum; °C) – the temperature at which the performance rate is maximized.
- R_max_ (maximal performance; µmol O_2_ cm^-2^ h^-1^) – the peak performance rate achieved at T_opt_.
- b(T_c_) – the performance rate (µmol O_2_ cm^-2^ h^-1^) measured at a reference temperature (defined here as 27°C, representing the ambient conditions during the TPC experiments).
- E_a_ (activation energy; eV) – the thermal sensitivity of the performance rate in the sub-optimal temperature range.
- E_h_ (deactivation energy; eV) – the rate of performance decline at supra-optimal temperatures.

### 2.8 | Statistical analyses

Each species was statistically evaluated separately. To assess the influence of historical bleaching phenotype and sampling year on physiological and thermal performance metrics, data were analyzed using linear mixed-effects models. To evaluate long-term recovery trajectories, physiological data (color score, biomass, symbiont density, P_max_) from 2023 and 2025 were integrated with previously published data from the 2019 bleaching event (data from October 2, 2019; Innis et al. 2021) and the 2021 recovery period from the same colony pairs (data from October 8, 2021; Brown et al. 2023). Prior to analysis, data were assessed for normality using quantile plots and the Shapiro-Wilk test. If normality assumptions were violated, data was either square-root or log-transformed to meet normality assumptions. For each species, linear mixed-effects models (nlme R package, v3.1-168; (Pinheiro et al. 2025)) with colony as a random intercept were fit for each metric with Year and Phenotype as fixed effects. The significance of the fixed effects and their interactions was determined via an ANOVA (type III) using the Anova function in the car R package (v3.1-5; (Fox and Weisberg 2019)). Significant effects were followed by pairwise comparison of estimated marginal means using the emmeans R package with Tukey HSD adjusted p values (v2.0.1; (Lenth and Piaskowski 2026)). Data analysis was done in R v4.5.0 (R Core Team 2024).

### 2.9 | Porites compressa symbiont community analysis

Previous work has found that *M. capitata* at PR13 host stable, phenotype-specific Symbiodiniaceae communities, typically dominated by *Cladocopium* (C31) in susceptible colonies and *Durusdinium glynnii* in resistant colonies (Cunning et al. 2016; Dilworth et al. 2021; Drury et al. 2022). However, it is unclear if such symbiont-phenotype associations exist for *P. compressa*. To this end, tissue biopsies (∼1–2 cm) for symbiont community analysis were collected from tagged resistant (n=8) and susceptible (n=8) *P. compressa* colonies in January 2025 using bone cutters. Fragments were immediately snap-frozen in liquid nitrogen and stored at -80°C until processing. Genomic DNA was extracted using a modified organic extraction protocol. Tissue was lysed in 1 mL of SDS buffer (0.1% SDS in 1x PBS) at 65°C for 1 hr, followed by a secondary incubation with 50 µL of proteinase K (20 mg/mL) at 55°C for 2–3 hrs. Phase separation was performed using phenol:chloroform:isoamyl alcohol (25:24:1), followed by a chloroform wash. DNA was precipitated via ethanol and CTAB (cetyltrimethylammonium bromide) treatments at 65°C. Purified gDNA pellets were washed in 70% ethanol, resuspended in ultrapure water, and quantified on a Nanodrop. The internal transcribed space 2 (ITS2) region of the Symbiodiniaceae ribosomal DNA was amplified using SYM_VAR forward and reverse primers modified with Illumina-compatible adapters (Hume et al. 2018). PCR was performed in 12.5 µL reactions containing 1 µL of template DNA and 11.5 µL of master mix (2X Phusion High-Fidelity PCR master mix (CAT M0531S; New England Biolabs) and 10 µM of each primer). Thermal cycling conditions included an initial denaturation at 98°C for 2 min; 34 cycles of 98°C for 10 s, 56°C for 30 s, and 72°C for 30 s; and a final extension at 72°C for 5 min. Amplification success and product size (∼400 bp) were confirmed by gel electrophoresis on a 2% agarose gel. PCR products were purified using the GeneJET PCR Purification Kit (CAT K0701; Thermo Fisher Scientific). Purified libraries were pooled, indexed, quality controlled and sequenced in a 2x300 bp configuration on an Illumina MiSeq at the Genomic Sequencing and Analysis Facility (GSAF) at the University of Texas at Austin, Center for Biomedical Research Support.

Raw FASTQ files were analyzed on the SymPortal remote server (https://symportal.org; (Hume et al. 2019).

Absolute abundance matrices were analyzed using the phyloseq R package (v1.52.0; (McMurdie and Holmes 2013). Relative abundances were calculated following removal of rare or low abundance variants, utilizing a >5% threshold per sample for ITS2 sequences (to capture dominant background variants) and a <1% threshold for ITS2 profiles (to capture distinct lineages) (Hume et al. 2019). Differences in symbiont community composition between phenotypes was evaluated with a PERMANOVA (Bray-Curtis dissimilarity) using the vegan package (v2.7-2; (Okasen et al. 2017)).

## 3 | Results

### 3.1 | Accumulation of thermal stress remained below the bleaching threshold in the six years following the 2019 MHW despite regular incursions above the MMM

Thermal profiles from March to October highlighted a clear contrast in accumulated heat stress between the 2019 heatwave and the subsequent recovery years (2021, 2023 and 2025; Figure 1A). The 2019 event represents the most recent marine heatwave in Kāneʻohe Bay, and was characterized by substantial heat accumulation (5.3 DHWs; Figure 1A, Figure S2), with sea surface temperatures exceeding the local bleaching threshold (28.3°C) for 22.3% of the March–October window (1,309 hours total) and remaining above the MMM (27.3°C) for 59.1% of the time (3,475 hours total; Figure 1B). In contrast, the 2021, 2023, and 2025 periods did not see accumulation of heat stress (Figure 1). Thermal stress accumulation was negligible, with zero DHWs recorded in 2021 and 2023 and a marginal accumulation of 0.19 DHWs observed prior to 2025 collections (Figure 1A, Figure S2). The frequency of exposure to temperatures above the bleaching threshold was also lower; temperatures exceeded the local bleaching threshold for 1.4% (83 hr; 2021), 0.1% (4 hr; 2023), and 2.2% (127 hr; 2025) of the season vs. 22.3% (1,309 hr) in 2019 (Figure 1B). While temperatures remained above the MMM for 14.6–26.2% of the period across these years (856–1,535 hr), this did not translate into accumulated degree heating weeks.

**Figure 1.**
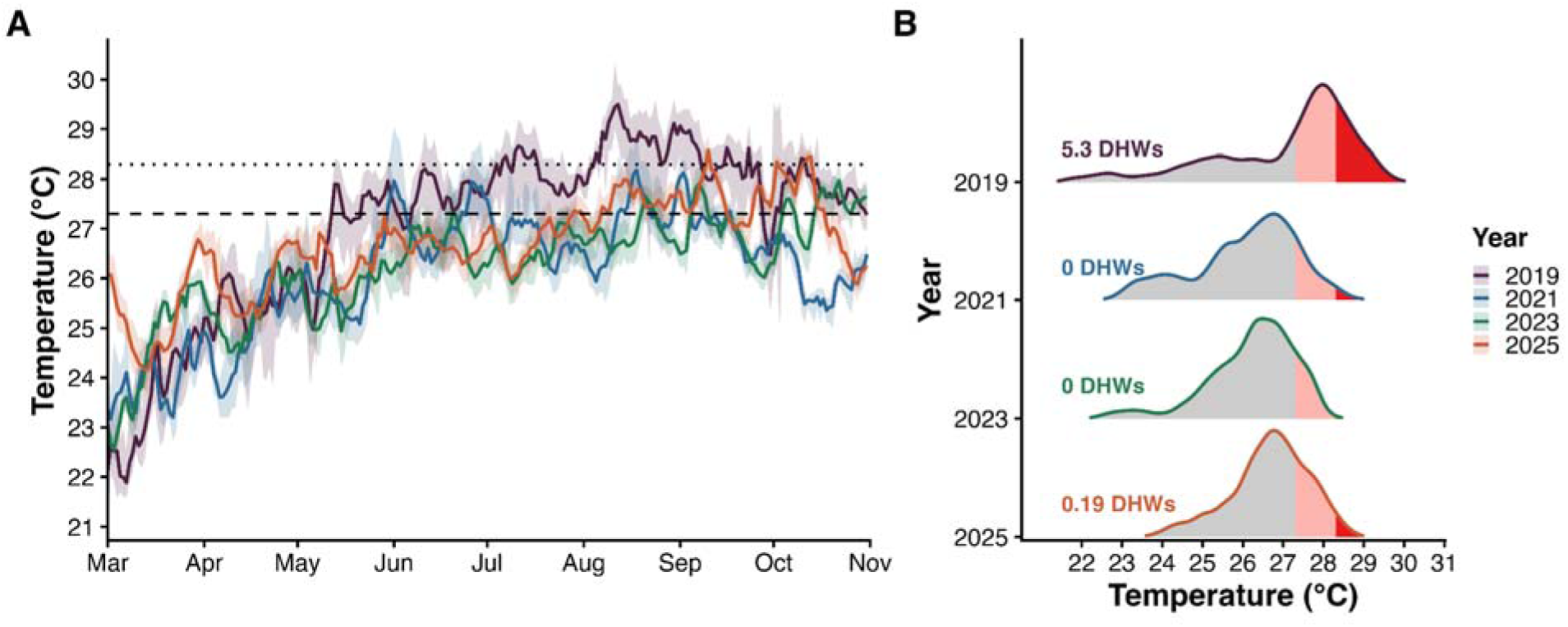
(A) Daily temperature profiles from March 1 through October 31 of 2019 (bleaching year), 2021 (non-bleaching year); 2023 (non-bleaching year), and 2025 (non-bleaching year). Solid lines represent the mean daily (24-hr) temperature, and shading represents the daily minimum and maximum temperatures. The dashed horizontal line indicates the maximum monthly mean (MMM; 27.3°C) of Kāneʻohe Bay, and the dotted horizontal line indicates the local coral bleaching threshold (MMM + 1°C; 28.3°C). (B) Probability density distributions of temperatures from March 1 through October 31 of 2019, 2021, 2023, and 2025. Shaded pink and red regions indicate the proportion of total time spent above the MMM (27.3°C) and the local coral bleaching threshold (28.3°C), respectively. The vertical height of each ridge represents the relative probability density, such that the total area under each curve equals one. The maximum amount of Degree Heating Weeks (DHWs) that each year accumulated during the March–October period is displayed to the left of the curve.

### 3.2 | Saturating light levels ensured metabolic rates were thermally limited during TPCs

Photosynthesis was saturated with respect to light at ∼500 µmol m^-2^ s^-1^ in September 2023 and ∼700 µmol m^-2^ s^-1^ in October 2025 across both species and phenotypes (Figure S3). Thus, photosynthesis and calcification rates during the TPC experiments were measured under saturating light levels (∼730 µmol m^-2^ s^-1^ in September 2023; ∼790 µmol m^-2^ s^-1^ in October 2025), ensuring that metabolic rates were limited by temperature rather than light availability.

### 3.3 | Long-term physiological recovery minimized initial phenotypic divergence in P. compressa but preserved it in M. capitata

Representative colony pairs are displayed in Figure S4. For *M. capitata*, the resistant phenotype maintained a consistently higher maximum photosynthesis (P_max_) than the susceptible phenotype (p=0.005; Table S4), with photosynthesis peaking in 2019 (Figure 2A). Overall, P_max_ was significantly influenced by year (*χ*^2^=8.415, p=0.038) and phenotype (*χ*^2^=10.386, p=0.001), but not their interaction (*χ*^2^=0.539, p=0.910; Figure 2A; Table S3). Following the 2019 peak, P_max_ decreased by 22.7% in 2021 (p=0.039), while 2023 and 2025 values only trended lower (12–13% reduction) without significantly differing from 2019 (Table S4). The resistant phenotype also maintained significantly higher symbiont density than the susceptible phenotype, averaging 68.5% higher across all years (p=0.012; Table S4). Symbiont density in *M. capitata* was significantly affected by year (*χ*^2^=39.494, p<0.001) and phenotype (*χ*^2^=7.766, p<0.001), but not their interaction (*χ*^2^=2.559, p=0.465; Figure 2B; Table S3). Across both phenotypes, symbiont density was at its minimum in 2019, rose by 203% by 2021 (p<0.001), and peaked in 2023 at 246% above 2019 levels (p<0.001; Table S4). By 2025, densities trended 35.8% lower than the 2023 peak (p=0.108) yet remained 122% higher than 2019 levels (p=0.003; Table S4). Biomass was significantly impacted by year (*χ*^2^=444.856, p<0.001), but not by phenotype (*χ*^2^=0.671, p=0.413) or their interaction (*χ*^2^=3.116, p=0.374; Figure 2C; Table S3). For both phenotypes, biomass increased by 128% from 2019 to 2021 (p<0.001) and peaked in 2023 (+217% relative to 2019), making 2023 significantly higher than all other years (Table S4). Biomass then dropped by approximately 40% in 2025 (p=0.006; Table S4). Similarly, color scores varied significantly by year (*χ*^2^=300.412, p<0.001), with no significant phenotype or interaction effects (p>0.4; Table S3; Figure 2D). Color scores were lowest during 2019 compared to all subsequent non-heatwave years (p<0.001; Table S4). Relative to 2019, color scores rose 49.8% in 2019, peaked in 2023 (+79.1%), and remained 61.4% higher in 2025 (Figure 2D). The 9.9% decline from 2023 to 2025 was significant (p<0.001), returning 2025 color scores to a level statistically similar to 2021 (p=0.074; Table S4).

**Figure 2.**
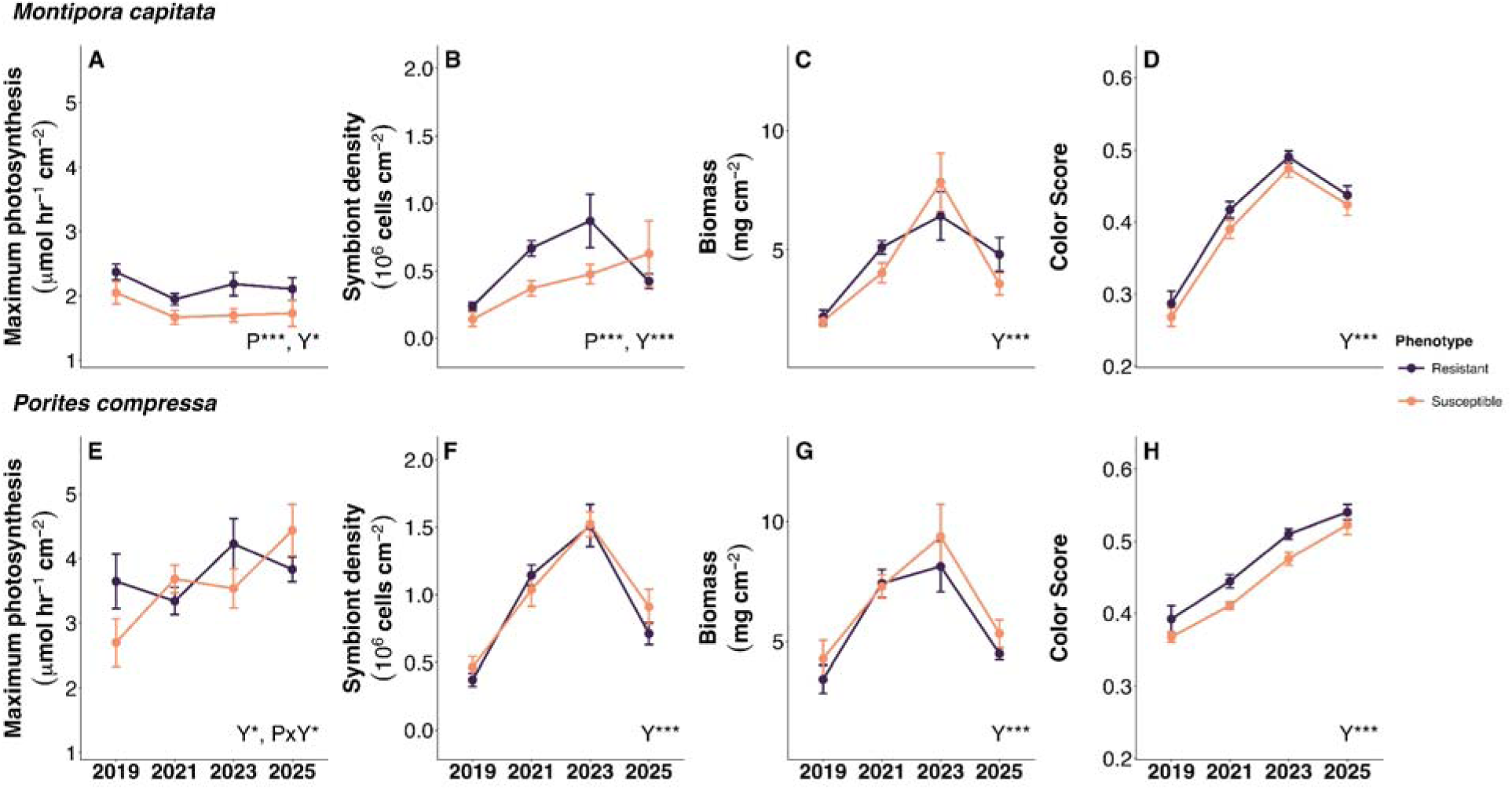
Comparative physiological metrics for *Montipora capitata* (top row; A–D) and *Porites compressa* (bottom row; E–H) across resistant (purple) and susceptible (orange) phenotypes in 2019, 2021, 2023, and 2025. Measured parameters include maximum photosynthetic rate (P_max_; A, E), symbiont density (B, F), biomass (C, G), and color score (D, H). Note that P_max_ presented here was derived from the photosynthesis-irradiance curves. Points and error bars represent the group means ± SEM. Statistical insets denote significant effects of year (Y), phenotype (P), or their interaction (YxP) based on linear mixed-effects models (* p<0.05; *** p<0.001; n.s. not significant).

For *P. compressa*, resistant colonies exhibited higher P_max_ than susceptible colonies in 2019 (p=0.049), though this phenotypic divergence disappeared in subsequent years (Table S6). Overall, P_max_ in *P. compressa* was significantly affected by year (*χ*^2^=9.840, p=0.019; Figure 2E) and a marginally significant interaction (*χ*^2^=8.092, p=0.054) but not by phenotype alone (*χ*^2^=0.422, p=0.516; Table S5; Figure 2E). The interaction was driven by the increase in P_max_ over time in susceptible colonies, culminating in a 64.4% increase by 2025 relative to 2019 (p=0.002; Table S6). Symbiont density in *P. compressa* varied significantly by year (*χ*^2^=104.948, p<0.001), but noy by phenotype (*χ*^2^=0.565, p=0.452) or their interaction (*χ*^2^=1.591, p=0.662; Table S5; Figure 2F). Across both phenotypes, symbiont density was lowest in 2019 and increased significantly in all subsequent years (p<0.05 for all comparisons, Table S6). Densities increased by 66.1% by 2021 (p<0.001), and peaked in 2023 (+94.6% relative to 2019; p<0.001; Table S6). Despite a 27.9% decline in 2025 (p<0.001), densities remained 40.3% higher than 2019 values (p<0.001; Table S6). Biomass followed a similar temporal pattern, shifting significantly by year (*χ*^2^=42.675, p<0.001), but not by phenotype (*χ*^2^=0.909, p=0.341) or their interaction (*χ*^2^=0.353, p=0.949; Table S5; Figure 2G). Both phenotypes demonstrated a sharp post-heatwave increase, rising 63.6% in 2021 (p<0.001) and 66.9% in 2023 (p<0.001; Table S6). By 2025, biomass dropped significantly from both 2021 (-22.2%, p=0.035) and 2023 (-23.8%, p=0.003) levels, returning to a state statistically similar to 2019 (p=0.105; Table S6). Color scores were shaped significantly by year (*χ*^2^=340.584, p<0.001) and phenotype (*χ*^2^=5.70, p=0.017), without a significant interaction (*χ*^2^=6.772, p=0.079; Table S5; Figure 2H). Resistant phenotypes maintained consistently higher color scores than susceptible ones across the study (p=0.021; Table S6). Scores were lowest in 2019 (p<0.001) and climbed continuously over time: +11.6% in 2021, +32.5% in 2023, and +46.0% in 2025. Every year-to-year step was statistically significant, including a final 10.2% increase from 2023 to 2025 (p<0.001; Table S6).

### 3.4 | Photosynthetic thermal optima remained fixed by phenotype in M. capitata but exhibited significant plastic shifts in P. compressa

Thermal performance curves measuring photosynthesis rates for *M. capitata* (Figure 3A, B) revealed significant interactions between phenotype and year for both the performance rate at 27°C (b(T_c_); *χ*^2^=4.718, p=0.030; Figure 4C) and activation energy (E_a_; *χ*^2^=5.580, p=0.018; Figure 4A; Table S7). Post-hoc analyses revealed that in 2023, resistant phenotypes exhibited marginally higher b(T_c_) (p=0.057) and significantly higher E_a_ (p<0.001) than susceptible colonies (Table S8). However, susceptible colonies showed significant increases in b(T_c_) (65.9%; p<0.001) and E_a_ (102%; p<0.001; Table S8) over time, eliminating phenotype differences in 2025 (b(T_c_) p=0.473; E_a_ p=0.792; Table S8). T_opt_ was also higher in resistant colonies (29.3°C) compared to susceptible colonies (28.4°C), representing a 0.9°C difference in thermal optima (*χ*^2^=7.33, p=0.007; Figure 4D; Table S8). Across both phenotypes, maximal photosynthetic performance (R_max_) significantly increased by 22.9% during the same period (*χ*^2^=6.85, p=0.009; Figure 4E; Table S7). No significant differences were observed in deactivation energy (E_h_) for *M. capitata* (Figure 4B; Table S8).

**Figure 3.**
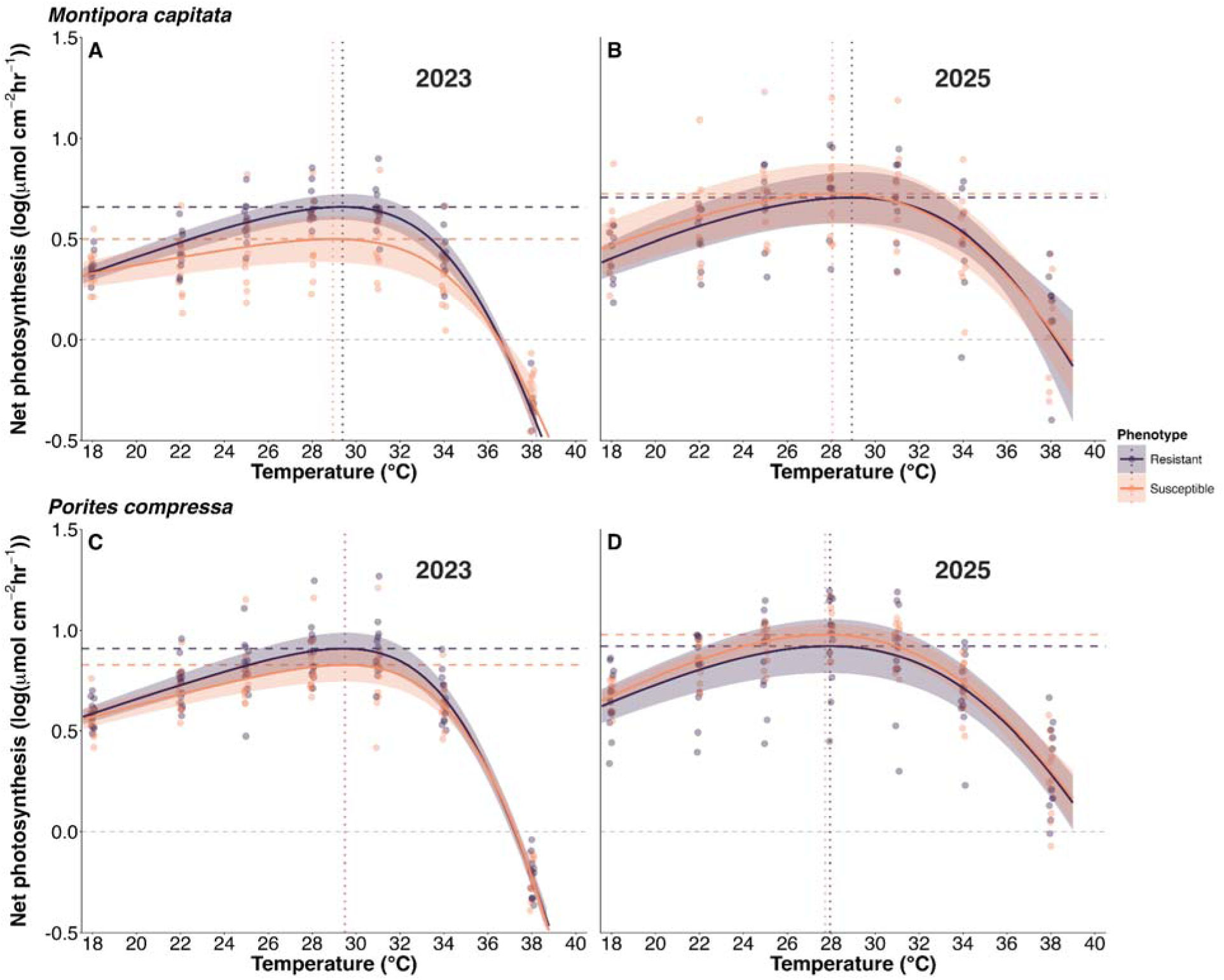
Thermal performance curves (TPCs) for net photosynthesis in two coral species. TPCs are shown for *Montipora capitata* (top row) in 2023 (A) and 2025 (B), and *Porites compressa* (bottom row) in 2023 (C) and 2025 (D) across resistant (purple) and susceptible (orange) phenotypes. Individual coral fragments are plotted as points, solid lines represent the best-fit curves, and shaded regions denote 95% confidence intervals. Horizontal dashed lines represent maximum photosynthesis (R_max_) and vertical dotted lines represent the thermal optimum (T_opt_) for each respective phenotype.

**Figure 4.**
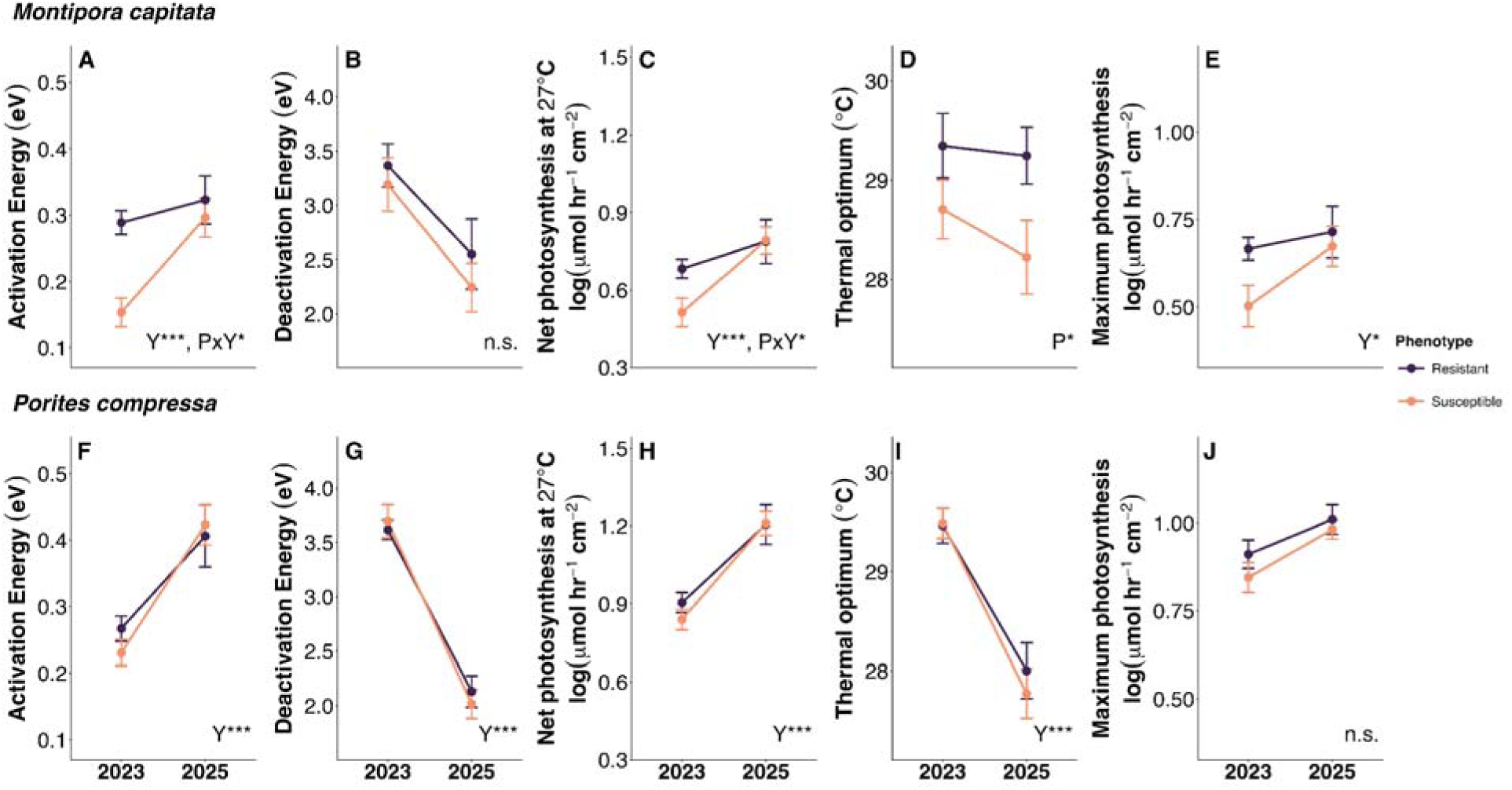
Thermal performance curve (TPC) metrics for net photosynthesis in two coral species. TPC metrics are shown for *Montipora capitata* (top row; A–E) and *Porites compressa* (bottom row; F–J) across resistant (purple) and susceptible (orange) phenotypes in 2023 and 2025. Extracted parameters include activation energy (E_a_; A, F), deactivation energy (E_h_; B, G), net photosynthesis at 27°C (b(T_c_); C, H), thermal optimum (T_opt_; D, I), and maximum photosynthesis (R_max_; E, J). Points and error bars represent the group means ± SEM. Statistical insets denote significant effects of year (Y), phenotype (P), or their interaction (YxP) based on linear mixed-effects models (* p<0.05; *** p<0.001; n.s. not significant).

Photosynthesis thermal performance metrics for *P. compressa* (Figure 3C, D) were primarily characterized by significant temporal shifts between years (Figure S4), with no significant differences observed between phenotypes or the interaction of phenotype and year (see Table S9). From 2023 to 2025, T_opt_ shifted downward by 1.6°C, decreasing from 29.5°C to 27.9°C (*χ*^2^=34.13, p<0.001; Figure 4I; Table S9). This shift in the peak was accompanied by significant changes in the slopes of the curve: E_a_ increased significantly (*χ*^2^=24.83, p<0.001; Figure 4F) and E_h_ decreased significantly (*χ*^2^=107.27, p<0.001; Figure 4G) over time (Table S9). Additionally, b(T_c_) increased by 35.3% from 2023 to 2025 (*χ*^2^=30.03, p<0.001; Figure 4H; Table S9). Despite modifications to the curve shape, R_max_ did not change significantly over the two year period (*χ*^2^=3.18, p=0.074; Figure 4J; Table S9).

### 3.5 | Calcification thermal performance remained stable in M. capiata but exhibited moderate changes in P. compressa over time

In contrast to the photosynthetic metrics, the thermal performance parameters for calcification in *M. capitata* (Figure 5A, B) did not differ between phenotypes and remained largely stable across years (Table S10). The only parameter impacted by an interaction between phenotype and year was E_a_ (*χ*^2^=3.84, p=0.050; Figure 6A; Table S10) yet post-hoc analyses revealed no significant pairwise differences between phenotypes or years (Table S11). While not statistically significant, qualitative trends in E_a_ suggested divergent trends between phenotypes: susceptible colonies showed an upward trend over time (p=0.106), while resistant colonies trended downward (p=0.351; Figure 6A; Table S11). All other parameters showed no significant response to the main effects or their interaction (p > 0.05; Figure 6B–E; Table S10).

**Figure 5.**
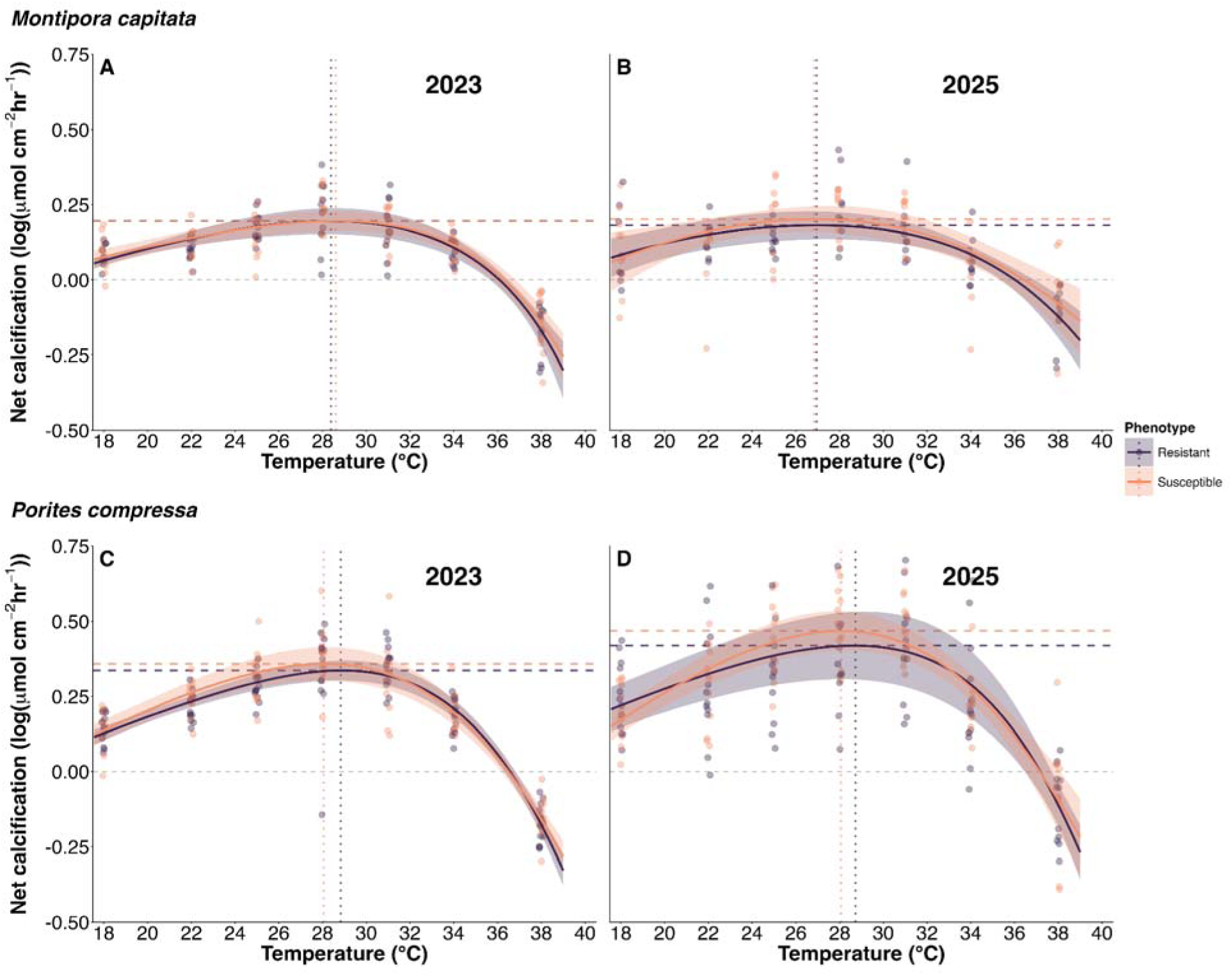
Thermal performance curves (TPCs) for net calcification in two coral species. TPCs are shown for *Montipora capitata* (top row) in 2023 (A) and 2025 (B), and *Porites compressa* (bottom row) in 2023 (C) and 2025 (D) across resistant (purple) and susceptible (orange) phenotypes. Individual coral fragments are plotted as points, solid lines represent the best-fit curves, and shaded regions denote 95% confidence intervals. Horizontal dashed lines represent maximum calcification (R_max_) and vertical dotted lines represent the thermal optimum (T_opt_) for each respective phenotype.

**Figure 6.**
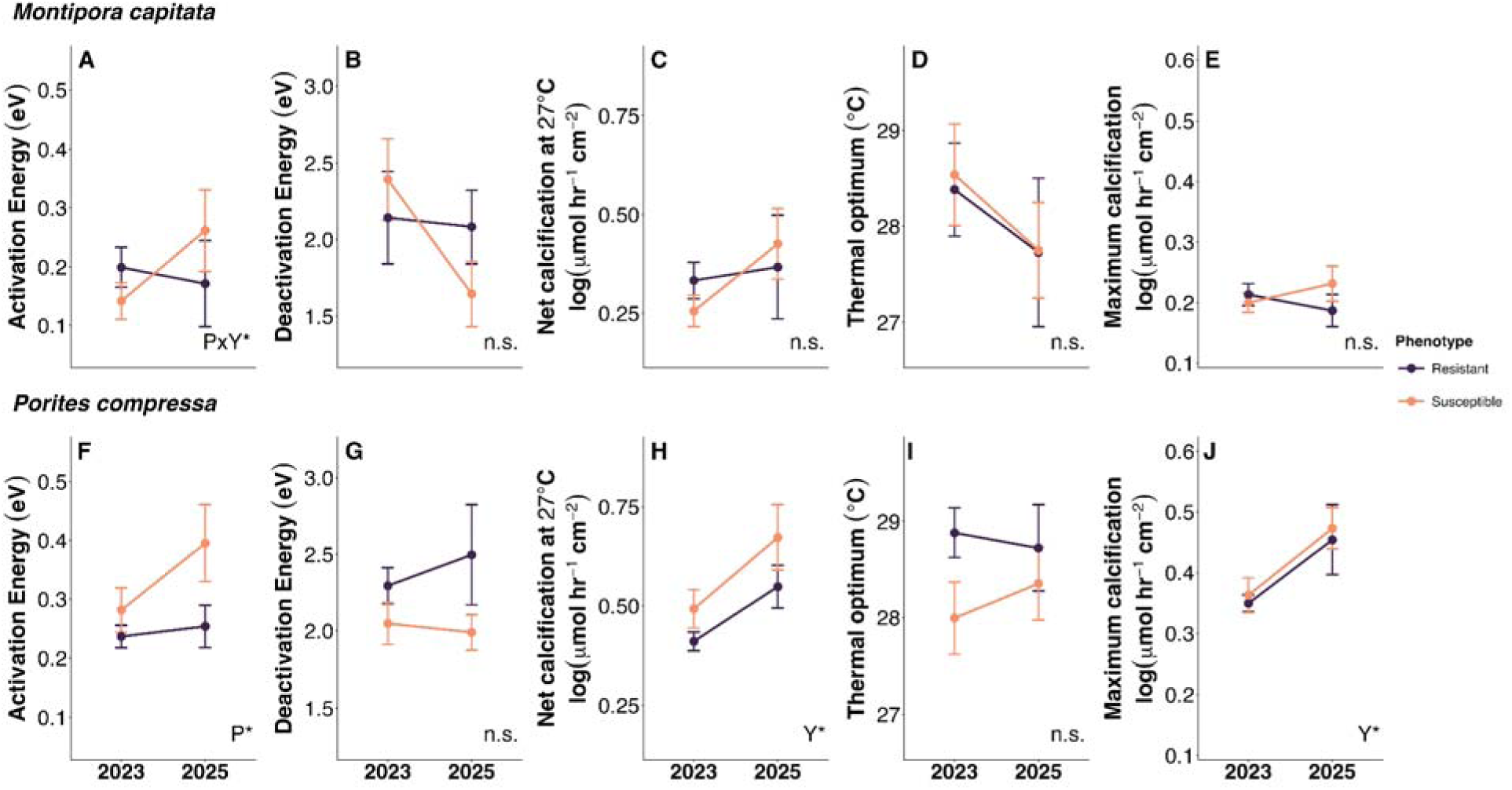
Thermal performance curve (TPC) metrics for net calcification in two coral species. TPC metrics are shown for *Montipora capitata* (top row; A–E) and *Porites compressa* (bottom row; F–J) across resistant (purple) and susceptible (orange) phenotypes in 2023 and 2025. Extracted parameters include activation energy (E_a_; A, F), deactivation energy (E_h_; B, G), net calcification at 27°C (b(T_c_); C, H), thermal optimum (T_opt_; D, I), and maximum calcification (R_max_; E, J). Points and error bars represent the group means ± SEM. Statistical insets denote significant effects of year (Y), phenotype (P), or their interaction (YxP) based on linear mixed-effects models (* p<0.05; *** p<0.001; n.s. not significant).

In contrast to *M. capitata*, certain thermal performance metrics for calcification in *P. compressa* (Figure 5C, D) were significantly influenced by year (Table S12). Regardless of phenotype, both the b(T_c_) (*χ*^2^=10.28, p=0.001; Figure 6H) and R_max_ (*χ*^2^=9.01, p=0.003; Figure 6J) increased significantly from 2023 to 2025, representing increases of 31.6% and 30.3%, respectively (Table S12). A marginally significant phenotypic effect was observed for E_a_ (*χ*^2^=3.56, p=0.059; Figure 6F), with susceptible colonies trending toward higher thermal sensitivity in the sub-optimal temperature range compared with resistant colonies (Table S12). No significant differences were observed for E_h_ or T_opt_ across years or phenotypes for calcification in *P. compressa* (p > 0.05; Figure 6G, I; Table S12).

### 3.6 | Cladocopium C15 maintains uniform dominance in P. compressa regardless of historical bleaching susceptibility

Symbiont community composition did not vary significantly between resistant and susceptible phenotypes of *P. compressa* (PERMANOVA p=0.587). NMDS ordination showed high overlap between phenotypes with a low stress value (0.087; Figure S5), indicating a robust representation of community structure. Relative abundance analysis demonstrated that both phenotypes were dominated by *Cladocopium* C15; specifically, C15 sequences consistently represented the majority of ITS2 reads across all colonies (Figure 7A), resulting in highly uniform *Cladocopium* C15 symbiont profiles (Figure 7B).

**Figure 7.**
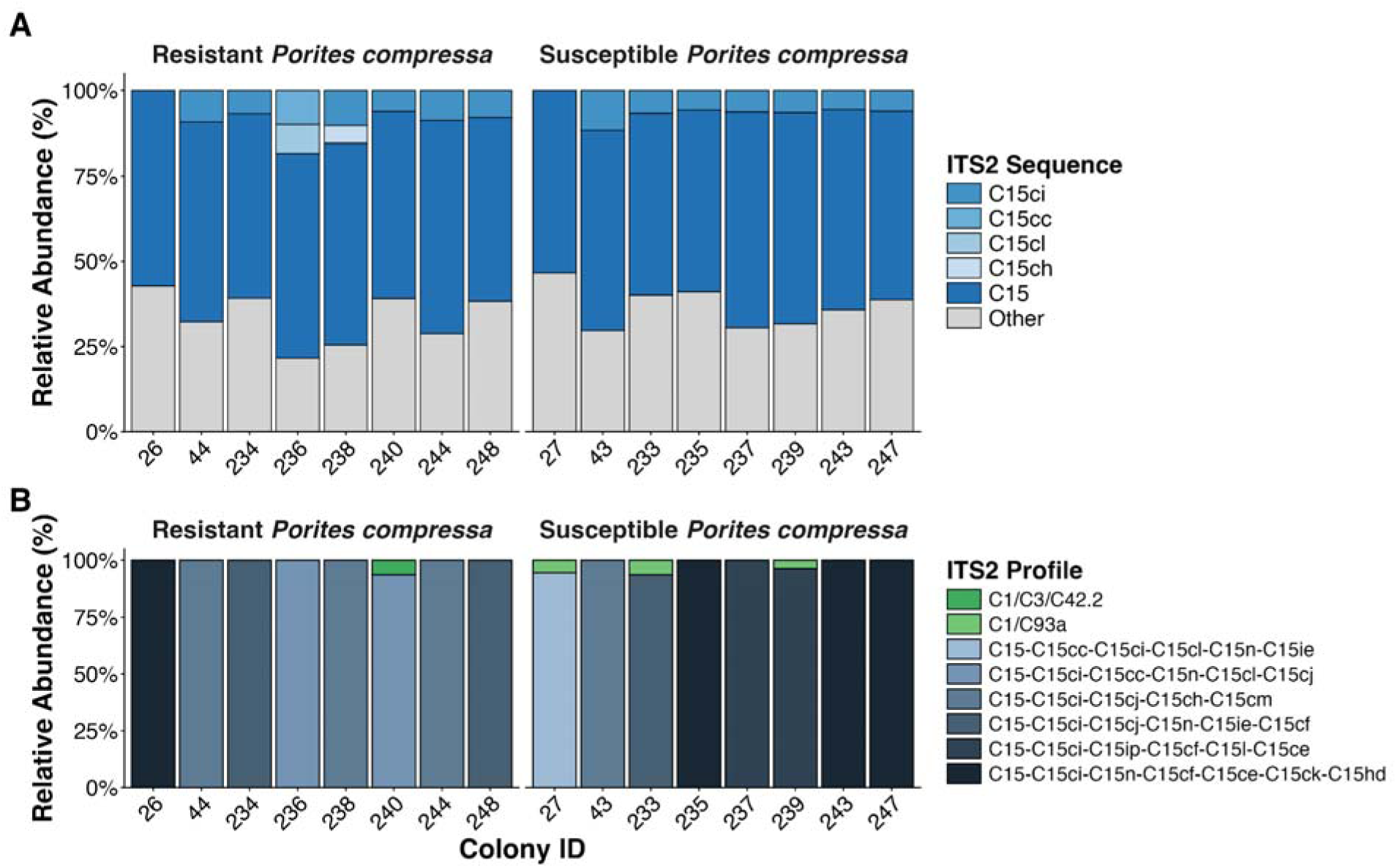
ITS2 sequence (A) and profile (B) composition in resistant and susceptible *Porites compressa*. (A) Relative abundance of dominant ITS2 sequences (>5% threshold per sample), with C15 variants and rare taxa (“Other”). (B) Relative abundance of SymPortal-defined ITS2 profiles (>1% threshold per sample). Samples are separated by phenotype (resistant and susceptible).

## 4 | Discussion

Corals in Kāneʻohe Bay continue to exhibit physiological legacies of heat stress for at least six years after a marine heatwave. We reveal that coral thermal performance is reshaped several years after bleaching but that recovery trajectories diverge by species, historical bleaching phenotype, and trait in ways that complicate straightforward assumptions of resilience. First, thermal optima (T_opt_) for photosynthesis in *M. capitata* remained fixed and distinct between phenotypes across time, demonstrating that their thermal limits may be constrained by symbiont identity (*Cladocopium* v. *Durusdinium*) even as other photosynthetic thermal performance traits converge. Second, *P. compressa* exhibited a striking 1.6°C downward recalibration of its photosynthetic thermal optimum toward ambient summer temperatures, indicating that elevated post-bleaching thermal tolerance may be transient in the prolonged absence of heat stress. Finally, photosynthesis and calcification recovered asynchronously within each species, with calcification in *P. compressa* still shifting measurably at six years post-bleaching—a timescale that, set against projections of near-annual bleaching by the end of the century (Spady et al. 2026) raises doubts about whether sufficient recovery windows will exist to restore full physiological function.

### 4.1 | Distinct thermal optima persist despite convergence of photosynthetic rates during recovery in M. capitata

In *M. capitata*, we observed a decoupling between the plasticity of photosynthetic performance and the stability of thermal optima (T_opt_). While basal photosynthetic rates (b(T_c_)) and metabolic acceleration (E_a_) converged between phenotypes over the two year study period, the 0.9°C gap in T_opt_ remained a fixed phenotypic marker, with resistant colonies maintaining a significantly higher thermal optima than susceptible conspecifics (29.3°C vs. 28.4°C, respectively). This suggests that the thermal optimum for photosynthesis is a trait in *M. capitata* that may not be subject to acclimatization but is instead more stable. Given projected increases in sea surface temperatures across Hawaiʻi over the next 50 years (1–2.5°C; (Ward et al. 2023)), this gap between bleaching-resistant and susceptible phenotypes will likely translate into long-term differences in bleaching frequency and survival under chronic ocean warming.

The divergence in thermal optima between bleaching phenotypes is likely rooted in the distinct symbiont communities each phenotype hosts. Bleaching-resistant *M. capitata* predominantly host the thermotolerant symbiont *Durusdinium glynnii*, while bleaching-susceptible colonies host primarily *Cladocopium* C31 symbionts (Cunning et al. 2016; Dilworth et al. 2021; Drury et al. 2022). These associations are remarkably stable in these colonies, with little evidence of post-bleaching symbiont ‘shuffling’, as bleached colonies typically recover their original dominant symbiont genera even after severe bleaching (Cunning et al. 2016; Dilworth et al. 2021). The higher thermal optimum for photosynthesis of *Durusdinium*-hosting (i.e., resistant) colonies likely reflects the intrinsic thermotolerant properties of this symbiont, including specialized lipid compositions to stabilize thylakoid membranes under heat stress (Rosset et al. 2019) and sustained carbon assimilation at temperatures that typically impair *Cladocopium* (Kemp et al. 2023; Liang et al. 2025). Notably, this phenotype-specific divergence was unique to *M. capitata* photosynthetic performance and was not observed in its calcification thermal performance. This mismatch likely reflects an energetic trade-off; thermotolerant *Durusdinium* translocate significantly less carbon to the *M. capitata* host than *Cladocopium* (Allen-Waller and Barott 2023), meaning that maintaining a higher photosynthetic thermal ceiling may not actually translate into enhanced host calcification. This discrepancy suggests that the photosynthetic thermal ceiling may reflect a symbiont-level constraint rather than a holobiont one. Whether this gap could eventually narrow through host-mediated processes, or represents an enduring feature of these distinct symbioses, remains an open question.

In contrast to the consistent differences of T_opt_, photosynthetic thermal performance converged between phenotypes over the study period. In 2023, resistant colonies had significantly higher basal performance (b(T_c_)) and a steeper metabolic acceleration (E_a_) than susceptible colonies, indicating incomplete recovery of susceptible colonies from the 2019 heatwave. This was further reflected in the lower biomass and symbiont densities in susceptible *M. capitata* colonies relative to resistant ones in 2023 (Brown, Lenz, et al. 2023). By 2025, phenotypes converged, with susceptible colonies recovering to performance levels comparable to resistant conspecifics. Interestingly, P_max_ in resistant corals remained higher across all time points, suggesting that the photosynthetic capacity in resistant, *Durusdinium*-hosting *M. capitata* may be sustained by a combination of host-mediated regulation of translocated carbon (Tremblay et al. 2012; Tremblay et al. 2014; Huffmyer et al. 2024), preferential maintenance of photosynthetic machinery (Scharfenstein et al. 2024; Liang et al. 2025), or other mechanisms. As such, four years following the marine heatwave (2023) captured *M. capitata* susceptible colonies in an active phase of photosynthetic recovery, while six years later (2025) represented a more complete restoration of the photosynthetic capacity.

For *M. capitata*, photosynthetic performance may be plastic and recoverable within limits, but the thermal ceiling may be constrained to some degree by symbiont identity, which can strongly modulate photosynthetic function and thermal tolerance (Jones and Berkelmans 2010; Turnham et al. 2023; Heit and Davy 2024). This contrast with *P. compressa*—in which photosynthetic T_opt_ showed plasticity across the same period in this study (Scharfenstein et al. 2024)—indicates that this rigidity is not universal but a product of the symbiotic relationship between the coral host and its symbiont communities. As marine heatwaves intensify and compress recovery windows, the distinction between which traits recover and what remains fixed may prove critical to predicting which coral species persist.

### 4.2 | Thermal recalibration in P. compressa may come at the expense of future heat tolerance

In contrast to *M. capitata*, photosynthetic TPC parameters in *P. compressa* exhibited no significant phenotypic or phenotype-by-year differences. This lack of differentiation is notable given the dramatically different bleaching responses observed in prior heatwaves (Matsuda et al. 2020). Instead, photosynthetic thermal performance was characterized by temporal shifts that impacted both phenotypes similarly. From 2023 to 2025, T_opt_ declined, E_a_ increased, E_h_ decreased and b(T_c_) increased, describing a thermal landscape that became narrower, taller, and peaked at a cooler temperature. This restructuring occurred alongside a decline in symbiont density and biomass from their 2023 peaks. These patterns suggest that *P. compressa* may be actively optimizing its photosynthetic efficiency for ambient conditions, even as it reduces its total symbiont load. This strategy aligns with broader patterns in coral physiology where lower symbiont densities reduce bleaching vulnerability (Cunning and Baker 2013) and shift holobiont photosynthesis to depend more on individual symbiont performance (Scheufen et al. 2017).

The 1.6°C downward shift in T_opt_ is a remarkably large physiological adjustment for *P. compressa* over just two years. The 2025 photosynthetic thermal optimum (27.9°C) falls within the temperature range that *P. compressa* typically experiences during non-heatwave summers in Kāneʻohe Bay (26–28°C), suggesting that photosynthesis was well-calibrated to the prevailing thermal environment by 2025. In contrast, the 2023 T_opt_ of 29.5°C sits above the local bleaching threshold (28.3°C) and may reflect an acclimation legacy or ‘environmental memory’ (Hackerott et al. 2021; Brown and Barott 2022) carried over from the 2019 heatwave. The shift observed between 2023 and 2025 indicates a potential limit to the duration of environmental memory as the coral recalibrates toward the prevailing ambient temperatures.

This recalibration raises an important and somewhat counterintuitive point: recovery toward a lower T_opt_ may signal a reduction in future heat tolerance. While the cooler 2025 T_opt_ represents *P. compressa* well-acclimated to present-day summer temperatures, the loss of the higher thermal ceiling maintained in 2023 leaves *P. compressa* less prepared to sustain metabolic performance during the next marine heatwave. This pattern is consistent with trade-offs documented in other corals, where optimization (acclimatization) for current temperatures comes at the cost of thermal breadth (Klepac and Barshis 2020; Roik et al. 2024). As marine heatwaves intensify and recovery windows shrink, the speed at which corals ‘reset’ their thermal limits—and potentially lose their protective legacy—will be a critical factor in forecasting reef vulnerability to recurrent thermal stressors (Putnam et al. 2017).

The uniformity of these temporal shifts across both *P. compressa* phenotypes may reflect the similarity of their symbiont communities. Colonies of both resistant and susceptible *P. compressa* phenotypes predominantly hosted *Cladocopium* C15, and this shared symbiont identity likely aligned with the convergent photosynthetic responses observed between phenotypes. However, because the symbiont community was only characterized six years post-heatwave in 2025, we cannot definitively rule out the possibility of symbiont shuffling during the intervening recovery period. Although both phenotypes share a broad taxonomic partnership with C15, these populations may be composed of distinct intra-type strains or low-abundance variants that exert different physiological effects on their host (Buerger et al. 2020; C. Wang et al. 2023). Given that these similar symbiont partnerships may have yielded divergent bleaching phenotypes in past heatwaves (Matsuda et al. 2020), the mechanisms driving differential bleaching thresholds may lie outside of symbiont species identity. Instead, these differences may be driven by host genetic or epigenetic variation (Torda et al. 2017; Eirin-Lopez and Putnam 2019; Drury 2020), strain-level physiological differences in symbiont populations (Buerger et al. 2020; C. Wang et al. 2023), microbiome composition (Barno et al. 2021; Núñez-Pons et al. 2023), or broader holobiont-level physiological differences (Barott et al. 2021; Wall et al. 2021). Disentangling these contributions will be important for understanding why differential bleaching responses occur in corals hosting the same species of symbionts.

### 4.3 | Photosynthetic and calcification thermal performance can recover asynchronously

Rather than recovering in parallel, photosynthesis and calcification TPC metrics followed distinct trajectories that differed by species and phenotype. In *M. capitata*, photosynthetic TPC parameters showed substantial phenotypic divergence in 2023 and subsequent convergence in 2025, reflecting an active metabolic recovery process still underway four years post-bleaching. Calcification TPC metrics, by contrast, remained largely stable across phenotypes and years, suggesting that *M. capitata* had already restored its calcification capacity to a functional steady state prior to our 2023 sampling, or alternatively, that this metric failed to recover to its historical pre-bleaching baseline. This lack of calcification divergence between phenotypes, even with a lower P_max_ in susceptible colonies and differences in TPC photosynthetic parameters in 2023, may be driven by symbiont identity once again. Although susceptible *M. capitata* colonies maintained a consistently lower P_max_ than their resistant counterparts, their *Cladocopium* symbionts are known to translocate a higher proportion of photosynthate to their hosts relative to colonies hosting *Durusdinium* (Allen-Waller and Barott 2023; C. Wang et al. 2023; Q. Wang et al. 2024). This enhanced carbon translocation efficiency likely allowed susceptible colonies to support calcification even at lower absolute P_max_ values. Alternatively, this calcification buffering may be explained by *M. capitata*’s heterotrophic flexibility, as this species can increase heterotrophic feeding following bleaching (Anthony and Fabricius 2000; Grottoli et al. 2006; Hughes and Grottoli 2013) to support energy-demanding processes like calcification. While we did not measure heterotrophy in our study, the stability of the *M. capitata* calcification TPCs, which were maintained even during periods of photosynthetic recovery, is consistent with this species buffering skeletal growth through alternative energy acquisition. By utilizing alternative energy pathways to sustain biomass and skeletal growth, *M. capitata* may be effectively insulating its calcification machinery from the prolonged recovery trajectory of its symbionts. However, more research is needed to fully validate this hypothesis.

In contrast to *M. capitata*, several calcification TPC parameters changed measurably over the study period in *P. compressa*. Because a pre-bleaching baseline is unavailable, these temporal shifts could be interpreted in two ways: either *P. compressa* had fully recovered by 2023 and its performance has continued to improve over time, or it was still actively undergoing a prolonged recovery six years post-heatwave. However, the consistent upward trends observed in both net calcification and ambient temperatures and maximal calcification rates suggests ongoing calcification recovery. This is consistent with previous work that has shown *Porites* spp. fail to recover calcification rates within the first eight months after bleaching (Schoepf et al. 2015) and may exhibit growth arrests lasting from one to four years after severe bleaching events (Cantin and Lough 2014; Gold and Palumbi 2018). Our data suggests that six years after the 2019 marine heatwave, *P. compressa* calcification is still undergoing measurable recovery. When contextualized with projections of annual bleaching by mid-century (Spady et al. 2026), this protracted timeline indicates that future disturbance intervals will outpace the capacity of these corals to recover, precluding the long-term stabilization of coral calcification. Together, these patterns indicate that although photosynthesis and calcification are linked energetically, they can recover on different timescales, and that those timescales may differ among coral species according to their physiological traits.

Across both species, we observed greater temporal change in photosynthetic TPCs than in calcification TPCs, a pattern that may reflect differences in the plasticity and cost of acclimating certain traits (Gattuso et al. 1999; Mass et al. 2007; Cohen and Dubinsky 2015). Photosynthetic performance can rapidly shift through changes in symbiont physiology (Middlebrook et al. 2012; Hoadley et al. 2019), photoprotective energy dissipation (Kana et al. 1997; Duanmu et al. 2017), and light harvesting adjustments (Duanmu et al. 2017). By contrast, calcification is a host-mediated process that requires coordinated ion transport (Davy et al. 2012; Barott et al. 2015), regulation of the calcifying fluid (Tambutté et al. 2011; Zoccola et al. 2015; Barott et al. 2020), and substantial energetic investment (Schoepf et al. 2013; Spalding et al. 2017), making it less amenable to rapid reconfiguration and therefore subject to greater constraints on the rate and magnitude of recovery. This aligns with the broader principle that the time required to adjust physiology and the energetic costs of that adjustment constrain how quickly and extensively organisms can shift the thermal performance of particular traits (Jurriaans and Hoogenboom 2020). Together, these patterns demonstrate that while photosynthesis and calcification are energetically linked, they recover on different timescales that is, in part, dictated by species-specific physiology, symbiont communities, and differences in thermal sensitivity among metabolic processes (Silbiger et al. 2019).

## Conclusions

By contextualizing physiological recovery within a decade-long tracking history of specific individual coral colonies, this study provides a rare and robust characterization of photosynthetic and calcification thermal performance in two dominant reef-building coral species during post-heatwave recovery. We demonstrate that coral thermal performance is not static but continues to be substantially reshaped in the absence of major heat stress events. Strikingly, we observed a downward recalibration of *P. compressa* photosynthetic T_opt_ toward ambient temperatures. This pronounced shift reflects acclimatization to prevailing conditions that may simultaneously reduce physiological preparedness and narrow safety margins ahead of future heat stress events. In *P. compressa*, no phenotypic differences in photosynthetic TPCs were detected, consistent with the uniform *Cladocopium* C15 symbiont community shared across both phenotypes but surprising given their contrasting historic bleaching susceptibility (Matsuda et al. 2020). However, given that these communities were only first characterized in early 2025, we cannot rule out the possibility of symbiont shuffling or disproportionate impacts on the holobiont from low-abundance strains. In *M. capitata*, resistant and susceptible phenotypes exhibited divergent photosynthesis TPC parameters in 2023 that converged by 2025, suggesting that physiological legacies of bleaching diminish with extended recovery. Nevertheless, persistent phenotypic divergence in *M. capitata* photosynthetic T_opt_ and P_max_ points to an enduring difference between phenotypes that may be partly constrained by symbiont identity (*Durusdinium* vs. *Cladocopium*). Calcification TPCs in *M. capitata* were largely stable across 2023 and 2025 and did not differ between bleaching phenotypes, even as photosynthetic TPC parameters continued to shift over that same period. In *P. compressa*, calcification TPCs changed over time, suggesting that this species had not yet fully restored its calcification capacity six years post-bleaching. Together, our findings underscore the importance of longitudinal studies and mechanistic frameworks such as TPCs for capturing the pace and trajectory of acclimatization. As recovery windows between bleaching events shorten, the speed at which corals can modify their thermal performance landscape will increasingly determine whether they can sustain critical metabolic functions over successive disturbances.

## Acknowledgements

We thank the staff of the Hawaiʻi Institute of Marine Biology and the Coral Resilience Lab for logistical and experimental support; Hollie Putnam for experimental support in 2023; Amelia Morgan and Julia Durian for color score analysis; Grace O’Brien for DNA extraction assistance; and Sarah Roderick for experimental support in 2025 and ITS2 analysis. Corals were collected under permits SAP-2024-24 (2023 experiment) and SAP-2026-21 (2025 experiment) issued by the state of Hawaiʻi’s Division of Aquatic Resources. This work was supported by the International Coral Reef Society (ICRS) Graduate Research Fellowship and the Penn Global Dissertation Grant to MPM, and the NSF CAREER award OCE 2237658 to KLB. The scientific results and conclusions, as well as any views or opinions expressed herein, are those of the author(s) and do not necessarily reflect the views of NOAA or the Department of Commerce.

## Data availability

All data and code required to reproduce the analyses and figures presented here are available in a public Github repository (https://github.com/JillAshey/PI_TPC) and will be archived on Zenodo following manuscript acceptance. Raw sequence data for ITS2 are available through the NCBI BioProject PRJNA1449003.

