## Supplemental Figures for "Decadal tracking reveals species-specific limits to coral thermal acclimatization"

**Supplemental materials**

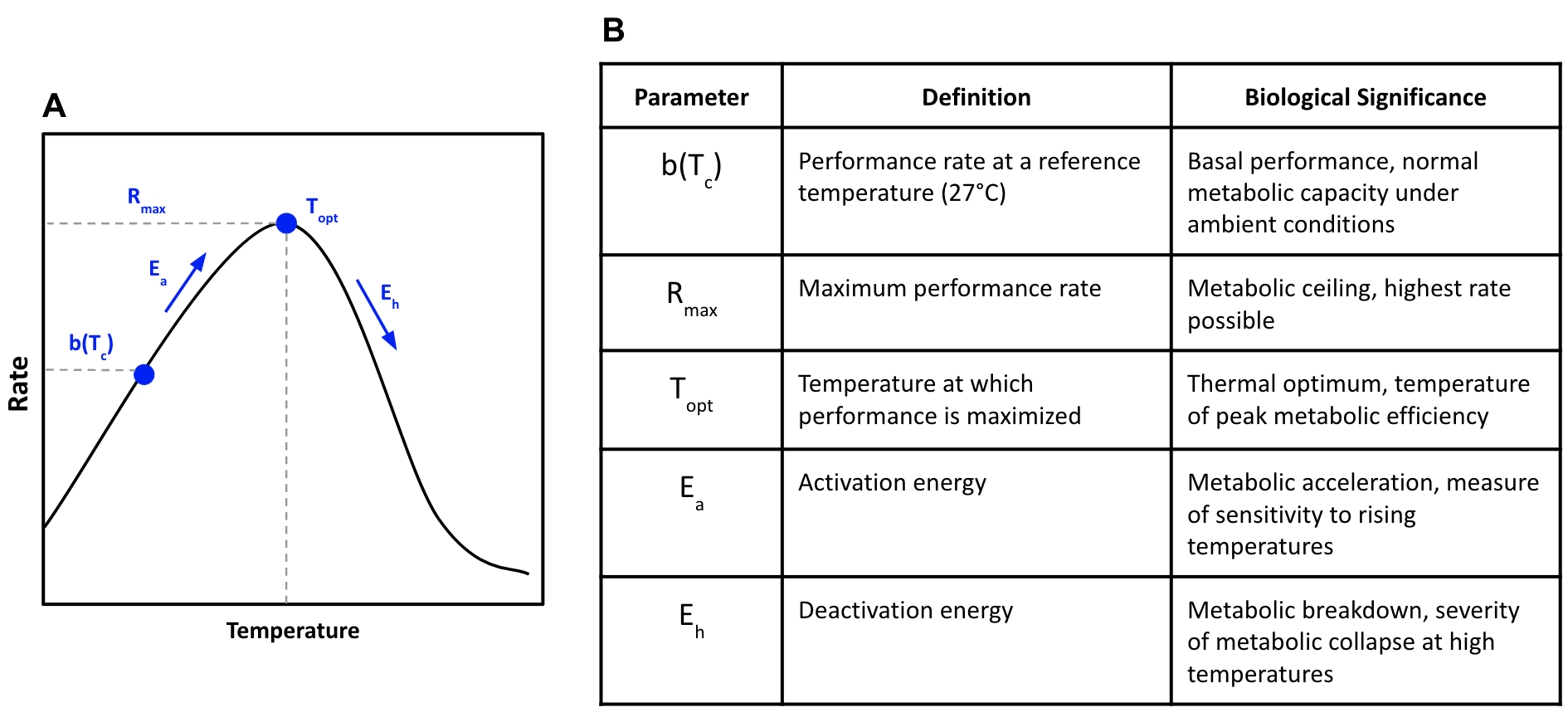

**Figure S1.** Parameters extracted from thermal performance curves. (A) A modeled thermal performance curve (Sharpe-Schoolfield) showing the relationship between performance rate and temperature. Metrics examined in this study are labeled on the graph. Adapted from Silbiger et al. 2019. (B) Definitions and biological significance of the thermodynamic and performance-based parameters derived from the model.

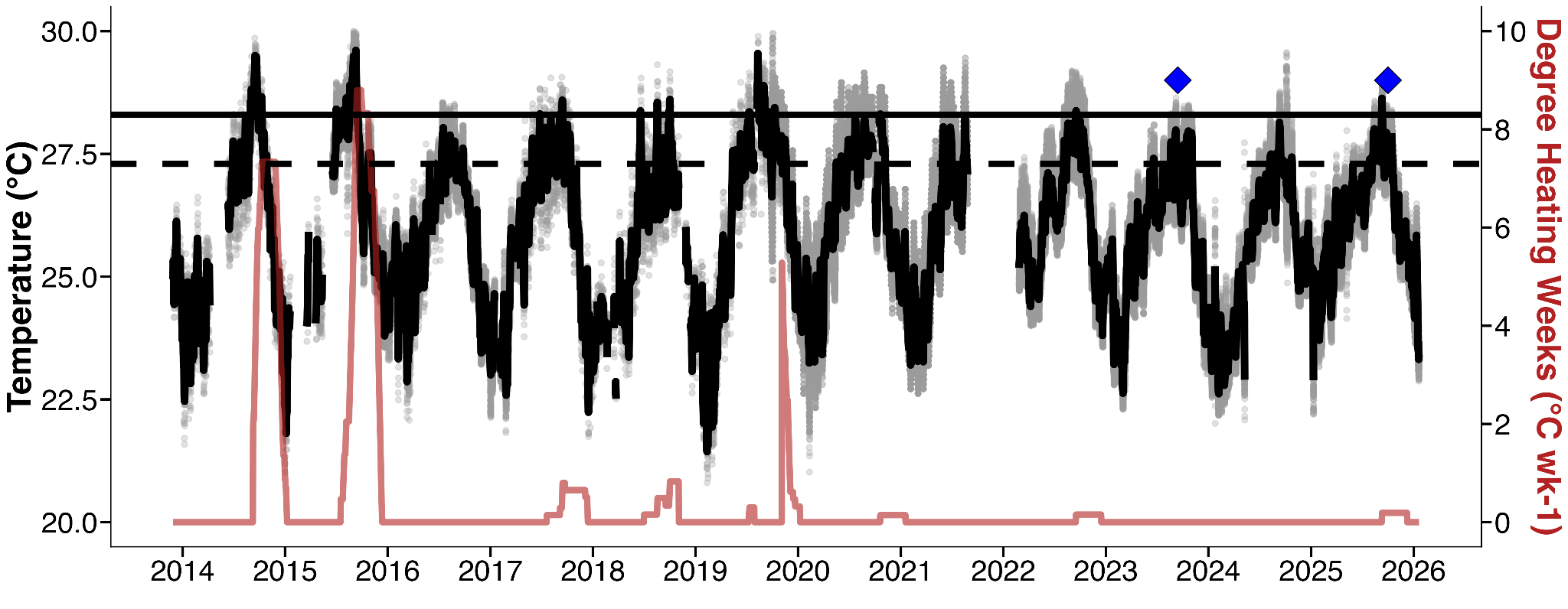

**Figure S2.** Temperature profiles and heat stress accumulation from 2014 to 2025 on Patch Reef 13 in Kāneʻohe Bay. Grey points indicate hourly measurements, with the black line representing the daily mean temperature. Heat stress accumulation was measured in degree heating weeks (DHWs; red line), calculated from mean daily temperatures. The dashed horizontal line indicates the maximum monthly mean (MMM; 27.3°C) of Kāneʻohe Bay, and the solid horizontal line indicates the local coral bleaching threshold (MMM + 1°C; 28.3°C). The blue triangles represent the thermal performance curve time points (September 2023 and October 2025) in this study.

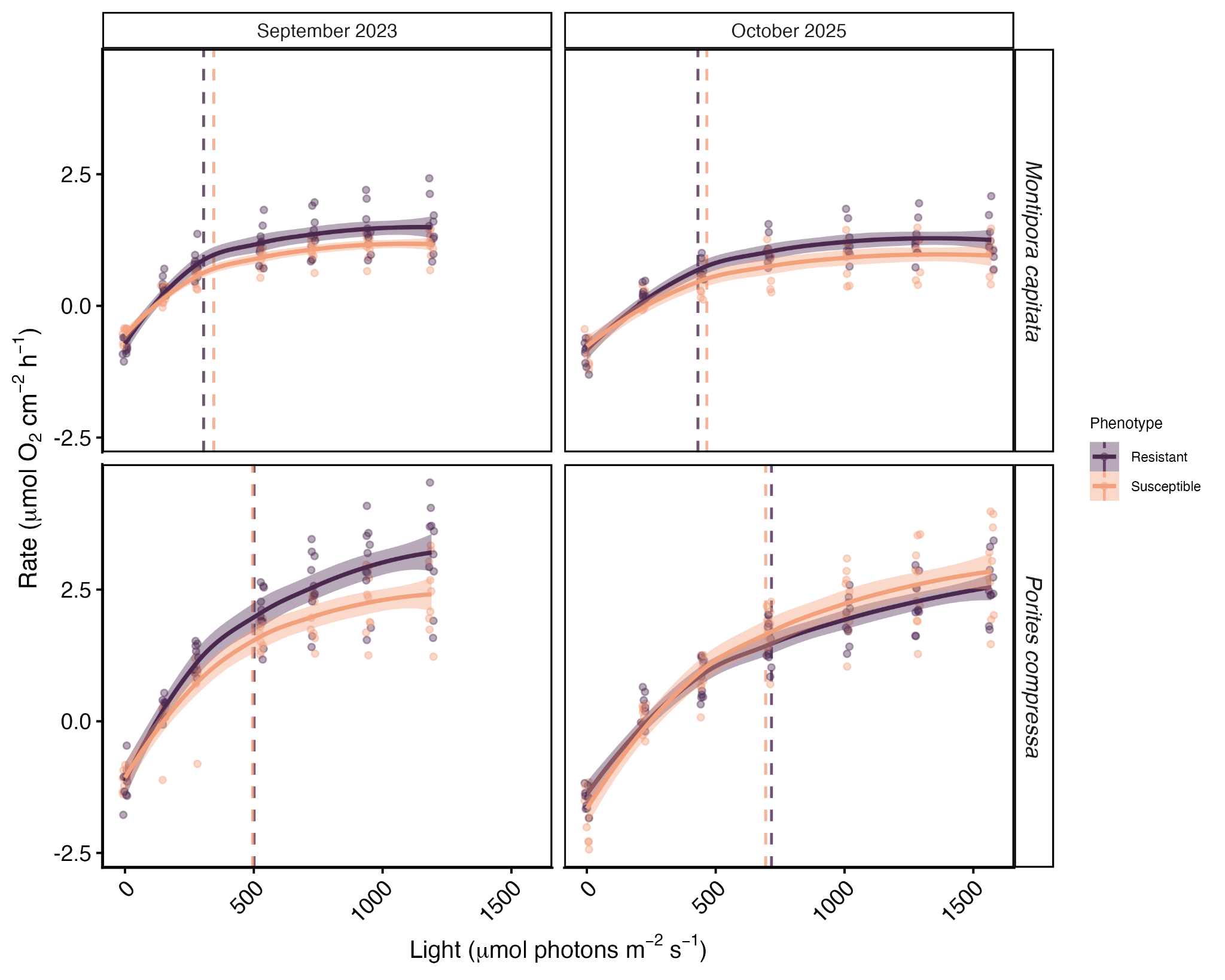

**Figure S3.** Photosynthesis-irradiance curves of bleaching-susceptible and bleaching-resistant *Montipora capitata* and *Porites compressa* in September 2023 (left panels) and October 2025 (right panels). Vertical dotted lines indicate saturating irradiance colored by phenotype.

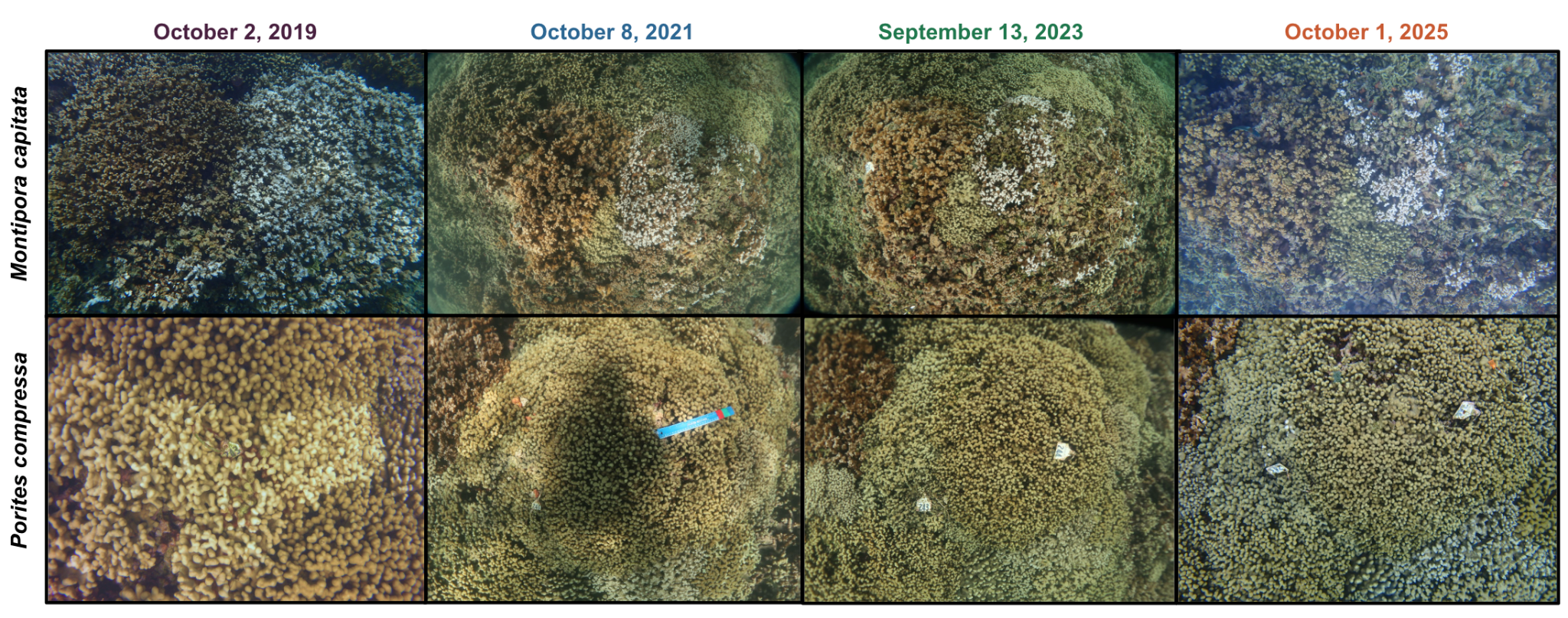

**Figure S4.** Representative images of Montipora capitata (top row) and Porites compressa (bottom row) pairs during the most recent marine heatwave (2019), +2 years post-heatwave recovery (2021), +4 years post-heatwave recovery (2023), and +6 years post-heatwave recovery (2025) in Kāneʻohe Bay, Hawaiʻi. Each row is the same two colonies over time.

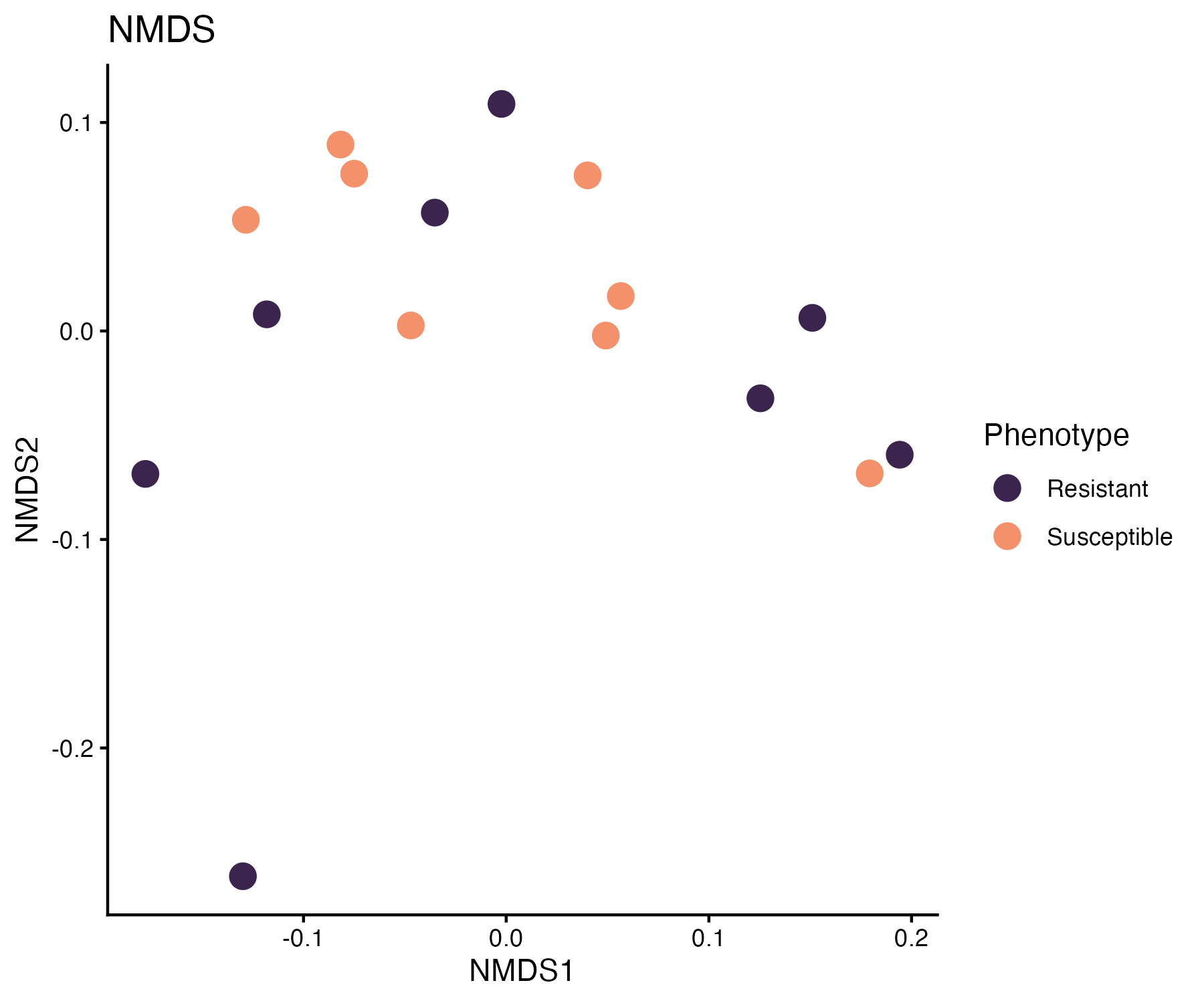

**Figure S5.** Non-metric multidimensional scaling (NMDS) of ITS2 (DIV) communities. The ordination is based on a Bray-Curtis distance matrix of filtered ITS2 sequence relative abundances (>1%). Each point represents a single colony.

**Table S1.** Coral sample sizes by date, species, and phenotype in September 2023 and October 2025.

| **Date** | **Species** | **Phenotype** | **Sample size** |
| --- | --- | --- | --- |
| September 2023 | *Montipora capitata* | Susceptible | 11 |
|  |  | Resistant | 11 |
|  | *Porites compressa* | Susceptible | 13 |
|  |  | Resistant | 13 |
| October 2025 | *Montipora capitata* | Susceptible | 10 |
|  |  | Resistant | 9 |
|  | *Porites compressa* | Susceptible | 12 |
|  |  | Resistant | 12 |

**Table S2.** Sample collection record for *Montipora capitata* and *Porites compressa* colonies for thermal performance curve experiments in September 2023 and October 2025. Colonies marked ‘Y’ indicate that the colony was sampled. Colonies marked ‘Dead’ means that the colony died by the 2025 experiment and was not sampled.

| **Colony ID** | **Species** | **Phenotype** | **TPC in 2023** | **TPC in 2025** |
| --- | --- | --- | --- | --- |
| 1 | *Montipora capitata* | Susceptible | Y | Y |
| 2 | *Montipora capitata* | Resistant | Y | Y |
| 3 | *Montipora capitata* | Susceptible | Y | Y |
| 4 | *Montipora capitata* | Resistant | Y | Y |
| 12 | *Montipora capitata* | Resistant | Y | Y |
| 19 | *Montipora capitata* | Susceptible | Y | Y |
| 20 | *Montipora capitata* | Resistant | Y | Y |
| 201 | *Montipora capitata* | Susceptible | Y | Y |
| 202 | *Montipora capitata* | Resistant | Y | Y |
| 203 | *Montipora capitata* | Susceptible | Y | Y |
| 204 | *Montipora capitata* | Resistant | Y | Y |
| 205 | *Montipora capitata* | Susceptible | Y | Y |
| 206 | *Montipora capitata* | Resistant | Y | Y |
| 209 | *Montipora capitata* | Susceptible | Y | Y |
| 210 | *Montipora capitata* | Resistant | Y | Y |
| 211 | *Montipora capitata* | Susceptible | Y | Y |
| 212 | *Montipora capitata* | Resistant | Y | Dead |
| 213 | *Montipora capitata* | Susceptible | Y | Y |
| 214 | *Montipora capitata* | Resistant | Y | Y |
| 219 | *Montipora capitata* | Susceptible | Y | Dead |
| 220 | *Montipora capitata* | Resistant | Y | Dead |
| 221 | *Montipora capitata* | Susceptible | Y | Y |
| 26 | *Porites compressa* | Resistant | Y | Y |
| 27 | *Porites compressa* | Susceptible | Y | Y |
| 28 | *Porites compressa* | Resistant | Y | Y |
| 29 | *Porites compressa* | Susceptible | Y | Y |
| 41 | *Porites compressa* | Susceptible | Y | Y |
| 42 | *Porites compressa* | Resistant | Y | Y |
| 43 | *Porites compressa* | Susceptible | Y | Y |
| 44 | *Porites compressa* | Resistant | Y | Y |
| 45 | *Porites compressa* | Susceptible | Y | Dead |
| 46 | *Porites compressa* | Resistant | Y | Dead |
| 225 | *Porites compressa* | Susceptible | Y | Y |
| 226 | *Porites compressa* | Resistant | Y | Y |
| 233 | *Porites compressa* | Susceptible | Y | Y |
| 234 | *Porites compressa* | Resistant | Y | Y |
| 235 | *Porites compressa* | Susceptible | Y | Y |
| 236 | *Porites compressa* | Resistant | Y | Y |
| 237 | *Porites compressa* | Susceptible | Y | Y |
| 238 | *Porites compressa* | Resistant | Y | Y |
| 239 | *Porites compressa* | Susceptible | Y | Y |
| 240 | *Porites compressa* | Resistant | Y | Y |
| 241 | *Porites compressa* | Susceptible | Y | Y |
| 242 | *Porites compressa* | Resistant | Y | Y |
| 243 | *Porites compressa* | Susceptible | Y | Y |
| 244 | *Porites compressa* | Resistant | Y | Y |
| 247 | *Porites compressa* | Susceptible | Y | Y |
| 248 | *Porites compressa* | Resistant | Y | Y |

**Table S3:** Full statistical results for physiological metrics in *Montipora capitata*. Bolding represents significant p-values.

| **Metric** | **Fixed effects** | **Chisq** | **Degrees of freedom** | **p-value (Pr > Chisq)** |
| --- | --- | --- | --- | --- |
| P_max_ | Phenotype | 10.386 | 1 | **0.001** |
|  | Year | 8.415 | 3 | **0.038** |
|  | Phenotype:  Year | 0.539 | 3 | 0.910 |
| Symbiont density | Phenotype | 7.766 | 1 | **<0.001** |
|  | Year | 39.494 | 3 | **<0.001** |
|  | Phenotype:Year | 2.559 | 3 | 0.465 |
| Biomass | Phenotype | 0.671 | 1 | 0.413 |
|  | Year | 444.856 | 3 | **<0.001** |
|  | Phenotype:Year | 3.116 | 3 | 0.374 |
| Color score | Phenotype | 0.458 | 1 | 0.499 |
|  | Year | 300.412 | 3 | **<0.001** |
|  | Phenotype:Year | 0.895 | 3 | 0.827 |

**Table S4:** Pairwise comparisons for significant effects of physiological metrics in *Montipora capitata*. Bolding represents significant p-values.

| **Metric** | **Contrast** | **df** | **t-ratio** | **p-value** |
| --- | --- | --- | --- | --- |
| P_max_ | 2019–2021 | 48.5 | 2.757 | **0.039** |
|  | 2019–2023 | 54.3 | 1.878 | 0.249 |
|  | 2019–2025 | 56.0 | 2.003 | 0.199 |
|  | 2021–2023 | 53.0 | -0.904 | 0.803 |
|  | 2021–2025 | 54.8 | -0.700 | 0.897 |
|  | 2023–2025 | 51.2 | 0.183 | 0.998 |
|  | Resistant–  Susceptile | 20.5 | 3.193 | **0.005** |
| Symbiont density | 2019–2021 | 54.9 | -5.104 | **<0.001** |
|  | 2019–2023 | 58.9 | -5.760 | **<0.001** |
|  | 2019–2025 | 62.8 | -3.687 | **0.003** |
|  | 2021–2023 | 53.0 | -0.668 | 0.909 |
|  | 2021–2025 | 56.3 | 1.534 | 0.424 |
|  | 2023–2025 | 50.8 | 2.317 | 0.108 |
|  | Resistant–  Susceptile | 19.7 | 2.767 | **0.012** |
| Biomass | 2019–2021 | 48.6 | -5.347 | **<0.001** |
|  | 2019–2023 | 53.6 | -7.775 | **<0.001** |
|  | 2019–2025 | 55.9 | -4.212 | **<0.001** |
|  | 2021–2023 | 52.9 | -2.161 | 0.148 |
|  | 2021–2025 | 55.4 | 1.150 | 0.661 |
|  | 2023–2025 | 49.5 | 3.460 | **0.006** |
| Color Score | 2019–2021 | 403 | -10.211 | **<0.001** |
|  | 2019–2023 | 416 | -17.151 | **<0.001** |
|  | 2019–2025 | 417 | -12.899 | **<0.001** |
|  | 2021–2023 | 407 | -6.421 | **<0.001** |
|  | 2021–2025 | 413 | -2.427 | 0.074 |
|  | 2023–2025 | 395 | 4.648 | **<0.001** |

**Table S5:** Full statistical results for physiological metrics in *Porites compressa*. Bolding represents significant p-values.

| **Metric** | **Fixed effects** | **Chisq** | **Degrees of freedom** | **p-value (Pr > Chisq)** |
| --- | --- | --- | --- | --- |
| P_max_ | Phenotype | 0.583 | 1 | 0.445 |
|  | Year | 9.822 | 3 | **0.020** |
|  | Phenotype:  Year | 8.092 | 3 | **0.054** |
| Symbiont density | Phenotype | 0.565 | 1 | 0.452 |
|  | Year | 104.948 | 3 | **<0.001** |
|  | Phenotype:Year | 1.591 | 3 | 0.662 |
| Biomass | Phenotype | 0.909 | 1 | 0.341 |
|  | Year | 42.675 | 3 | **<0.001** |
|  | Phenotype:Year | 0.353 | 3 | 0.949 |
| Color score | Phenotype | 5.700 | 1 | **0.017** |
|  | Year | 340.584 | 3 | **<0.001** |
|  | Phenotype:Year | 7.201 | 3 | 0.066 |

**Table S6:** Pairwise comparisons for significant effects of physiological metrics in *Porites compressa*. Bolding represents significant p-values.

| **Metric** | **Contrast** | **df** | **t-ratio** | **p-value** |
| --- | --- | --- | --- | --- |
| P_max_ | 2019–2021 | 47.9 | -0.946 | 0.780 |
|  | 2019–2023 | 55.6 | -2.129 | 0.157 |
|  | 2019–2025 | 54.6 | -2.922 | **0.025** |
|  | 2021–2023 | 55.1 | -1.018 | 0.739 |
|  | 2021–2025 | 53.8 | -1.731 | 0.318 |
|  | 2023–2025 | 45.3 | -0.766 | 0.869 |
|  | 2019–2021 in Resistant | 49.0 | 0.535 | 0.950 |
|  | 2019–2023 in Resistant | 55.6 | -1.230 | 0.611 |
|  | 2019–2025 in Resistant | 55.6 | -0.387 | 0.980 |
|  | 2021–2023 in Resistant | 53.8 | -1.689 | 0.339 |
|  | 2021–2025 in Resistant | 53.8 | -0.905 | 0.803 |
|  | 2023–2025 in Resistant | 44.5 | 0.872 | 0.819 |
|  | 2019–2021 in Susceptible | 46.8 | -1.904 | 0.240 |
|  | 2019–2023 in Susceptible | 55.6 | -1.781 | 0.293 |
|  | 2019–2025 in Susceptible | 53.6 | -3.797 | **0.002** |
|  | 2021–2023 in Susceptible | 56.6 | 0.226 | 0.996 |
|  | 2021–2025 in Susceptible | 53.8 | -1.543 | 0.419 |
|  | 2023–2025 in Susceptible | 46.2 | -1.914 | 0.236 |
|  | Resistant–  Susceptible in 2019 | 64.0 | 2.011 | **0.049** |
|  | Resistant–  Susceptible in 2021 | 64.0 | -0.515 | 0.608 |
|  | Resistant–  Susceptible in 2023 | 64.0 | 1.460 | 0.149 |
|  | Resistant–  Susceptible in 2025 | 64.0 | -1.326 | 0.189 |
| Symbiont density | 2019–2021 | 65.2 | -5.934 | **<0.001** |
|  | 2019–2023 | 67.7 | -9.499 | **<0.001** |
|  | 2019–2025 | 68.3 | -4.124 | **<0.001** |
|  | 2021–2023 | 59.0 | -2.909 | **0.026** |
|  | 2021–2025 | 61.2 | 2.681 | **0.045** |
|  | 2023–2025 | 53.3 | 6.711 | **<0.001** |
| Biomass | 2019–2021 | 56.8 | -4.754 | **<0.001** |
|  | 2019–2023 | 61.6 | -5.815 | **<0.001** |
|  | 2019–2025 | 63.4 | -2.315 | 0.105 |
|  | 2021–2023 | 60.1 | -0.251 | 0.994 |
|  | 2021–2025 | 62.2 | 2.788 | **0.035** |
|  | 2023–2025 | 52.2 | 3.676 | **0.003** |
| Color Score | 2019–2021 | 349 | -4.039 | **0.004** |
|  | 2019–2023 | 358 | -12.731 | **<0.001** |
|  | 2019–2025 | 359 | -17.454 | **<0.001** |
|  | 2021–2023 | 353 | -7.749 | **<0.001** |
|  | 2021–2025 | 354 | -12.368 | **<0.001** |
|  | 2023–2025 | 349 | -6.633 | **<0.001** |
|  | Resistant–  Susceptible | 98.4 | 2.340 | **0.021** |

**Table S7:** Full statistical results for metrics extracted from the photosynthesis thermal performance curves in *Montipora capitata*. Bolding represents significant p-values.

| **TPC Metric** | **Fixed effects** | **Chisq** | **Degrees of freedom** | **p-value (Pr > Chisq)** |
| --- | --- | --- | --- | --- |
| b(T_c_) | Phenotype | 0.483 | 1 | 0.487 |
|  | Year | 16.079 | 1 | **<0.001** |
|  | Phenotype:  Year | 4.718 | 1 | **0.030** |
| E_a_ | Phenotype | 7.629 | 1 | **0.005** |
|  | Year | 13.383 | 1 | **<0.001** |
|  | Phenotype:Year | 5.580 | 1 | **0.018** |
| E_h_ | Phenotype | 0.049 | 1 | 0.884 |
|  | Year | 0.021 | 1 | 0.842 |
|  | Phenotype:Year | 0.039 | 1 | 0.843 |
| T_opt_ | Phenotype | 7.333 | 1 | **0.007** |
|  | Year | 1.015 | 1 | 0.313 |
|  | Phenotype:Year | 0.489 | 1 | 0.484 |
| R_max_ | Phenotype | 1.205 | 1 | 0.272 |
|  | Year | 6.847 | 1 | **0.009** |
|  | Phenotype:Year | 2.926 | 1 | 0.087 |

**Table S8:** Post-hoc results for significant interaction effects of metrics extracted from the photosynthesis thermal performance curves in *Montipora capitata*. Bolding represents significant p-values.

| **TPC Metric** | **Interaction** | **Context** | **Contrast** | **df** | **t-ratio** | **p-value** |
| --- | --- | --- | --- | --- | --- | --- |
| b(T_c_) | Phenotype \| Year | 2023 | Resistant– Susceptible | 33.5 | 1.971 | **0.057** |
|  |  | 2025 | Resistant –Susceptible | 35.1 | -0.725 | 0.473 |
|  |  | Resistant | 2023 – 2025 | 19.0 | -1.239 | 0.231 |
|  |  | Susceptible | 2023 – 2025 | 17.2 | -4.534 | **<0.001** |
| E_a_ | Phenotype \| Year | 2023 | Resistant– Susceptible | 36.0 | 3.833 | **<0.001** |
|  |  | 2025 | Resistant –Susceptible | 36.0 | 0.266 | 0.792 |
|  |  | Resistant | 2023 – 2025 | 20.0 | -0.881 | 0.389 |
|  |  | Susceptible | 2023 – 2025 | 17.8 | -4.381 | **<0.001** |

**Table S9:** Full statistical results for metrics extracted from the photosynthesis thermal performance curves in *Porites compressa*. Bolding represents significant p-values.

| **TPC Metric** | **Fixed effects** | **Chisq** | **Degrees of freedom** | **p-value (Pr > Chisq)** |
| --- | --- | --- | --- | --- |
| b(T_c_) | Phenotype | 0.001 | 1 | 0.973 |
|  | Year | 30.028 | 1 | **<0.001** |
|  | Phenotype:  Year | 1.872 | 1 | 0.171 |
| E_a_ | Phenotype | 0.006 | 1 | 0.937 |
|  | Year | 24.826 | 1 | **<0.001** |
|  | Phenotype:Year | 1.916 | 1 | 0.166 |
| E_h_ | Phenotype | 0.080 | 1 | 0.777 |
|  | Year | 107.269 | 1 | **<0.001** |
|  | Phenotype:Year | 0.449 | 1 | 0.505 |
| T_opt_ | Phenotype | 0.063 | 1 | 0.802 |
|  | Year | 34.132 | 1 | **<0.001** |
|  | Phenotype:Year | 0.109 | 1 | 0.741 |
| R_max_ | Phenotype | 0.015 | 1 | 0.903 |
|  | Year | 3.183 | 1 | 0.074 |
|  | Phenotype:Year | 2.072 | 1 | 0.150 |

**Table S10:** Full statistical results for metrics extracted from the calcification thermal performance curves in *Montipora capitata*. Bolding represents significant p-values.

| **TPC Metric** | **Fixed effects** | **Chisq** | **Degrees of freedom** | **p-value (Pr > Chisq)** |
| --- | --- | --- | --- | --- |
| b(T_c_) | Phenotype | 0.012 | 1 | 0.914 |
|  | Year | 0.758 | 1 | 0.384 |
|  | Phenotype:  Year | 3.427 | 1 | 0.064 |
| E_a_ | Phenotype | 0.071 | 1 | 0.789 |
|  | Year | 0.381 | 1 | 0.527 |
|  | Phenotype:Year | 3.835 | 1 | **0.050** |
| E_h_ | Phenotype | 0.101 | 1 | 0.751 |
|  | Year | 1.734 | 1 | 0.188 |
|  | Phenotype:Year | 1.909 | 1 | 0.167 |
| T_opt_ | Phenotype | 0.004 | 1 | 0.953 |
|  | Year | 2.113 | 1 | 0.146 |
|  | Phenotype:Year | 0.081 | 1 | 0.776 |
| R_max_ | Phenotype | 0.429 | 1 | 0.513 |
|  | Year | 0.004 | 1 | 0.948 |
|  | Phenotype:Year | 2.050 | 1 | 0.152 |

**Table S11:** Post-hoc results for significant interaction effects of metrics extracted from the calcification thermal performance curves in *Montipora capitata*. Bolding represents significant p-values.

| **TPC Metric** | **Interaction** | **Context** | **Contrast** | **df** | **t-ratio** | **p-value** |
| --- | --- | --- | --- | --- | --- | --- |
| E_a_ | Phenotype \| Year | 2023 | Resistant– Susceptible | 30.6 | 1.143 | 0.262 |
|  |  | 2025 | Resistant –Susceptible | 32.0 | -1.263 | 0.216 |
|  |  | Resistant | 2023 – 2025 | 15.3 | 0.961 | 0.351 |
|  |  | Susceptible | 2023 – 2025 | 16.5 | -1.710 | 0.106 |

**Table S12:** Full statistical results for metrics extracted from the calcification thermal performance curves in *Porites compressa*. Bolding represents significant p-values.

| **TPC Metric** | **Fixed effects** | **Chisq** | **Degrees of freedom** | **p-value (Pr > Chisq)** |
| --- | --- | --- | --- | --- |
| b(T_c_) | Phenotype | 2.027 | 1 | 0.155 |
|  | Year | 10.277 | 1 | **0.001** |
|  | Phenotype:  Year | 0.381 | 1 | 0.537 |
| E_a_ | Phenotype | 3.563 | 1 | **0.059** |
|  | Year | 2.138 | 1 | 0.144 |
|  | Phenotype:Year | 1.637 | 1 | 0.201 |
| E_h_ | Phenotype | 2.996 | 1 | 0.083 |
|  | Year | 0.011 | 1 | 0.916 |
|  | Phenotype:Year | 0.025 | 1 | 0.874 |
| T_opt_ | Phenotype | 2.213 | 1 | 0.137 |
|  | Year | 0.094 | 1 | 0.759 |
|  | Phenotype:Year | 0.492 | 1 | 0.483 |
| R_max_ | Phenotype | 0.204 | 1 | 0.652 |
|  | Year | 9.007 | 1 | **0.003** |
|  | Phenotype:Year | 0.006 | 1 | 0.937 |
